# A mechanistically novel peptide agonist of the IL-7 receptor that addresses limitations of IL-7 cytokine therapy

**DOI:** 10.1101/2023.05.24.542196

**Authors:** William J. Dower, Angie Inkyung Park, Alice V. Bakker, Steven E. Cwirla, Praechompoo Pongtornpipat, Blake M. Williams, Prarthana Joshi, Bryan A. Baxter, Michael C. Needels, Ronald W. Barrett

## Abstract

Interleukin (IL)-7 is broadly active on T-cell populations, and modified versions have been clinically evaluated for a variety of therapeutic applications, including cancer, lymphopenia and infectious diseases; and found to be relatively well-tolerated and biologically active. Here we describe novel IL-7R agonists that are unrelated in structure to IL-7, bind to the receptor subunits differently from IL-7, but closely emulate IL-7 biology. The small size, low structural complexity, and the natural amino acid composition of the pharmacologically active peptide MDK1472 allows facile incorporation into protein structures, such as the IgG2-Fc fusion MDK-703. This molecule possesses properties potentially better suited to therapeutic applications than native IL-7 or its derivatives. We compared these compounds with IL-7 for immune cell selectivity, induction of IL-7R signaling, receptor-mediated internalization, proliferation, and generation of immune cell phenotypes in human and non-human primate (NHP) peripheral blood cells in vitro; and found them to be similar in biological activity to IL-7.

In cynomolgus macaques, MDK-703 exhibits a circulating half-life of 46 hr, and produces sustained T-cell expansion characteristic of IL-7 treatment. In the huCD34^+^-engrafted NSG mouse model of the human immune system, MDK-703 induces an immune cell profile very similar to that generated by IL-7-derived compounds; including the pronounced expansion of memory T-cells, particularly the population of stem-like memory T-cells (Tscm), which may be important for anti-tumor activities reported with IL-7 treatment.

Clinical administration of IL-7 and modified variants has been reported to induce anti-drug antibodies (ADAs), including IL-7 neutralizing antibodies. The novel peptide agonist reported here scores very low in predicted immunogenicity, and because the peptide lacks sequence similarity with IL-7, the problematic immunogenic neutralization of endogenous cytokine should not occur.

## Introduction

Interleukin-7 receptor (IL-7R) signaling is essential to establishing and sustaining immuno-competence throughout life. Most phases of T-cell development, and the continuing maintenance of T-cell homeostasis and functional responsiveness of the lymphoid system, depend on the availability of IL-7 and well-regulated control of its receptor expression. The absence of T-cells in patients with IL-7Rα mutations results in severe combined immune deficiency, revealing the essential role of the IL-7 system in healthy immune function [1, 2].

Unlike other cytokines regulating T-cell biology, IL-7 is produced primarily by non-hematopoietic tissues, among the most active of which is the stroma of lymphoid organs, including thymus and lymph nodes [3]. Upon secretion, most IL-7 is sequestered locally through binding to epithelia and ECM [4-6], and is detectable in the peripheral circulation of healthy humans at concentrations less than 10 pg/mL. Under normal conditions, T-cells experience limiting levels of IL-7 and must compete for this cytokine. While IL-7 levels are low and constant, IL-7Rα expression fluctuates, mediating cell selectivity and sensitivity [7]. Under conditions of T-lymphocyte depletion (lymphopenia caused by a variety of conditions such as viral infection, sepsis, cancer, and therapeutic interventions), the reduced number of T-cells internalize less IL-7, thereby elevating both local and circulating levels to provide proliferative stimulus and potential reversal of the lymphopenic condition [8].

The IL-7 receptor belongs to the family of interleukin receptors that share the common gamma subunit (γc). γc is ubiquitously expressed on immune cells, while private cytokine-specific α or β subunits are uniquely distributed among immune cell types, conferring the pattern of cell selectivity for each of the common gamma cytokines. Regulation of surface density of the IL-7Rα subunit is a principal determinant of cell sensitivity to IL-7 stimulation; and IL-7 exposure modulates the surface density of IL-7Rα at two levels. First, upon IL-7 binding and initiation of signaling, the activated receptor complexes are rapidly internalized, and IL-7Rα is directed to late endosomes for degradation; second, IL-7R signaling down regulates transcription of IL-7Rα. Besides the immediate effect on the cell surface level of IL-7Rα, internalization of the signaling complex also depletes or “consumes” IL-7. The resulting decline in IL-7 and IL-7R signaling relieves transcriptional suppression, allowing IL-7Rα expression and cell sensitivity to IL-7 to rise. This fluctuation of IL-7 availability and IL-7Rα expression is intrinsic to the feedback regulation of T-lymphocyte homeostasis. Recognition of this oscillating dynamic led to the concept of IL-7 sharing (or competition) among T-cells that depend on the limited supply of IL-7 for survival. This shared access to IL-7 seems key to maintaining healthy steady-state T-cell populations - the immune homeostasis feature that characterizes IL-7 biology [9-11].

Among the cells most consistently responsive to IL-7 are naive and memory T-cells; both are long-lived lymphocytes requiring periodic IL-7R stimulation. Throughout life, immuno-competence depends on a naive T-cell population retaining proliferative and differentiation potential, and IL-7 drives thymic generation, peripheral proliferation and survival, and T-cell receptor (TCR) diversification to maintain a robust population of potential effectors. Following immune response, IL-7 provides ongoing support of a select population of effectors that have differentiated into memory cells [12].

Recently, IL-7R agonists have been shown to drive the generation of a “stem-like” population of early memory/effector cells believed important for tumor suppression. IL-7 diversifies the TCR repertoire by inducing the expansion of less differentiated but more proliferative T naïve (T_N_) and stem cell-like memory T (T_SCM_) subpopulations. T_SCM_ are a subset of memory T-lymphocytes with the ability to rapidly self-renew and the capacity to reconstitute a spectrum of memory and effector T-cell subsets. T_SCM_ may be necessary for the control of persistent pathologies as antigen-driven effector cells undergo functional exhaustion and replicative senescence, requiring continuous replenishment by less differentiated T-cell subsets [13-15].

IL-7, through its influence on many lymphocyte classes, exerts broad influence on immune readiness and response, and long-term memory capability, which suggests a variety of immune-therapeutic applications for exogenous IL-7R agonists. This clinical potential is supported by many animal studies evaluating effects of recombinant forms of IL-7 in models of lymphopenia, infectious disease and sepsis, and cancer [11, 16-18]. In human studies, recombinant IL-7 (non-glycosylated and glycosylated), and a pharmacokinetic (PK)-enhanced Fc-fusion of IL-7 (efineptakin alfa) have been clinically evaluated for use in lymphopenia, septic shock, infectious disease, enhancement of response to vaccination, lymphopenia associated with COVID-19, autoimmune diseases, chronic inflammatory diseases, and cancer. These IL-7R agonists appear to be relatively well-tolerated and biologically active in the clinic [16, 19-26].

Among the various indications, there is particularly strong interest in the use of IL-7R agonists for treating cancer. Recombinant human (rh)IL-7 and modified forms, administered to mice, NHP, and humans have been shown to induce widespread T-cell proliferation, increase T-cell numbers, modulate peripheral T-cell subsets, and increase TCR repertoire diversity. In oncology applications, these biological attributes, alone or in combination with other immune-based therapies exhibit anti-tumor effects in mice and may enhance the rate, depth, and durability of clinical responses. IL-7R agonists alone or in combination with checkpoint inhibitors have been reported to increase the number and function of anti-tumor T lymphocytes in peripheral blood of several cancer patient populations (e.g., checkpoint inhibitor [CPI]-naïve relapsed/refractory microsatellite stable colorectal cancer [MSS-CRC], CPI-naïve pancreatic cancer, and newly diagnosed high-grade gliomas after chemoradiation) [13, 27, 28].

The most intensively studied IL-7R agonist now in development is the PK-enhanced IL-7 efineptakin alfa, previously shown to suppress tumor growth in preclinical murine tumor models, both as monotherapy and in combination with a PD-1 CPI. Efineptakin alfa in the clinic (as monotherapy and in combination with PD-(L)1 inhibitors) is relatively well-tolerated and biologically active, achieving considerable T-cell expansion with limited T_REG_ expansion.

Efineptakin alfa has achieved clinical responses (partial and complete) when combined with CPI in immunologically cold tumor types and tumors that are relapsed/refractory to prior treatments. In various clinical studies of efineptakin alfa in combination with a CPI, increased T-cell proliferation in the periphery, and objective responses have been observed in immunologically cold tumor types (microsatellite stable colorectal cancer, pancreatic ductal adenocarcinoma) and tumor types that are relapsed/refractory to prior treatments (triple negative breast cancer, non-small cell lung cancer, and small cell lung cancer)[29-33]. Here we describe novel agonists of the human IL-7 receptor, discovered from randomly assembled peptides and unrelated to the sequence of IL-7; and we compare these surrogate agonists with IL-7 in stimulating relevant T-cell subsets. The compounds discussed are a synthetic 42 amino acid peptide, MDK1472, and MDK-703, a recombinant fusion of the amino acid sequence of MDK1472 to an IgG2 Fc domain to extend half-life *in vivo*. We show these compounds to be full agonists in inducing immune cell phenotypes that closely emulate those generated by natural IL-7. The small size and low structural complexity of the pharmacologically active peptide component allows facile incorporation into protein structures, such as therapeutic antibodies, to combine with IL-7 agonism to create agents with new biological properties. These agonist molecules score very low in predicted immunogenicity analysis; and the lack of sequence similarity with IL-7 avoids the possibility of immunogenic neutralization of the endogenous cytokine if an ADA response should occur.

## Results

### Peptide ligands of IL-7 receptor subunits identified and assembled into receptor agonists

The IL-7R is a heterodimer consisting of a cytokine-specific alpha chain (IL-7Rα, CD127) and the common gamma chain (γc, CD132), the latter shared by receptors of six cytokines belonging to the γc family. As is typical of hematopoietic cytokine receptors, IL-7R signaling is induced by engagement of protein agonists with both receptor subunits, either recruiting individual subunits into an oligomeric complex, or re-arranging pre-assembled subunits to correctly position the subunit-associated janus kinases (JAKs) to initiate a signaling cascade. To create molecules that emulate this agonistic mechanism, we employed an iterative process of peptide ligand identification for each of the two receptor subunits, ligand affinity enhancement, combinatorial assembly of ligand pairs into heterodimers, and functional testing of heterodimeric peptides to identify those exhibiting IL-7R agonist activity.

To identify collections of ligands binding to either human IL-7Rα or γc receptor subunits, we constructed libraries of peptides expressed on filamentous phage (each >10^10^ clones per library) comprised of randomly assembled peptides, with or without positions containing fixed amino acids chosen to induce secondary structure; and these were screened for binding to the target subunit extracellular domains (ECDs), as we have previously described [34, 35]. Screening campaigns yielded distinct families of peptide ligands specific for each of the subunits. Each family was defined by a set of many sequence-related peptides, none of which exhibited sequence similarity to IL-7. Representative peptides, closely aligned with the consensus sequence of each family, were chemically synthesized and tested for receptor subunit binding. These initial hits bound only to the subunit from which they were recovered, typically with low affinities, often weaker than 1 μM.

Affinity maturation of these initial families produced ligands with improved affinities [34]; and combinations of these ligands (one each for IL-7Rα and γc) were joined into heterodimers in a variety of orientations (i.e., joined by N-termini, C-termini, or N-to-C termini), with several connection chemistries, for testing as IL-7R agonists in cell-based assays. The lead peptide agonist for detailed characterization, as described here, is a fusion of peptides MDK1248 and MDK1188, joined with a short intervening linker to form the single-chain peptide MDK1472. The sequences of the individual peptide ligands and the resulting single-chain peptide agonist are shown in Fig 1. The peptides were analyzed for sequence similarity to IL-7, and none was found. MDK1472 exhibited IL-7R agonist activity as a synthetic peptide. MDK1472 is relatively stable when incubated *in vitro* with human or cynomolgus plasma at 37°C (t_1/2_ = 60 hrs and 104 hrs, respectively, S1 Fig), but its modest size (4836 Da) leaves it vulnerable to rapid renal clearance and limited circulating half-life *in vivo*. To improve pharmacokinetic performance, we constructed a fusion of MDK1472 to the C-termini of both chains of human IgG2 Fc-domain, a compound designated MDK-703 (Fig 1C); and we have shown this molecule to closely emulate the pharmacological properties of the synthetic peptide MDK1472.

**Fig 1.**
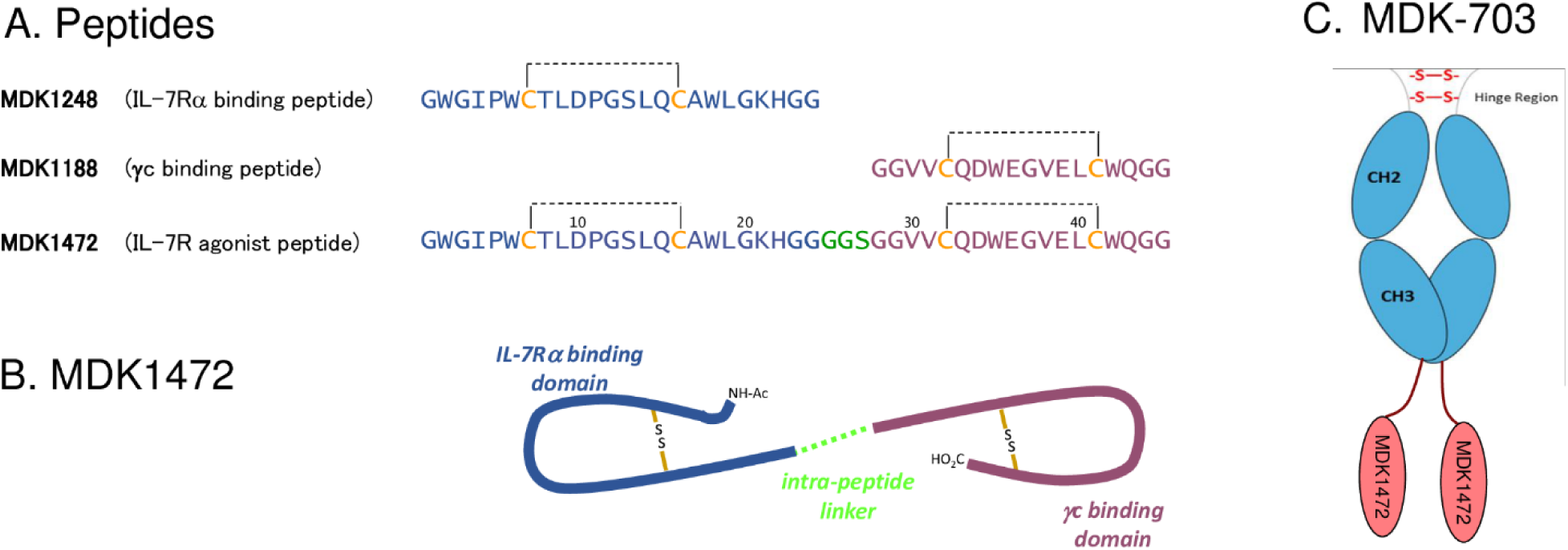
Sequences and schematic structures of the compounds. (A) Amino acid sequences of MDK1248, MDK1188, and MDK1472 with intrachain disulfide bonds shown as dashed lines. Schematic representations of (B) peptide MDK1472 and (C) IgG2Fc-1472 fusion MDK-703.

The stochastic process of discovery of MDK1472 was unbiased by information from naturally occurring proteins, and led to the discovery of ligands with unique amino acid sequences, unrelated to IL-7 or other proteins in the human protein sequence database. In therapeutic applications, such non-self peptides have the potential for generating an immune response in patients, leading to ADAs that may neutralize the activity of the drug. To estimate the likelihood of this possibility, a segment of the Fc-fusion MDK-703, including the sequence beginning 9 residues upstream of the linker that connects the Fc-domain to MDK1472 and continuing to the C-terminus of the MDK1472 sequence was evaluated by an algorithmic predictive screen for the presence of human class II (HLR “DR”) restricted HLA epitopes (EpiVax, Inc., EpiMatrix™ system) [36, 37]. This screen analyzes the set of nested 9-mers covering the entire surveyed sequence (including linker sequences) for predicted recognition by the 12 major MHC haplotypes. The low global “EpiMatrix™” score and regional analysis for clusters of potential HLA-reactive segments suggest limited potential for immunogenicity (score of -21.70); MDK-703 lacks evidence of putative human HLA DR epitope clusters and was chosen as the lead candidate to advance into preclinical development. Because active peptide constituents of MDK1472 and MDK-703 are completely novel, we have undertaken a broad comparison of the functional properties of these compounds with human IL-7.

### Binding and agonist properties of MDK1472 and MDK-703

#### IL-7R subunit binding of ligands and agonists

Binding affinities of peptide MDK1472 and its constituent subunit-binding fragments, MDK1248 (IL-7Rα) and MDK1188 (γc), and of the Fc fusion MDK-703 to human and cynomolgus subunits were determined by competition binding enzyme-linked immunosorbent assay (ELISA). The IC_50_s of MDK1248 binding human IL-7Rα ECD, and MDK1188 binding human γc ECD are 150 and 24 nM, respectively (Figs 2A and 2B), and are similar to those of the heterodimeric peptide MDK1472 (340 and 57 nM); while MDK-703 (presenting the heterodimeric peptide on each Fc chain) binds IL-7Rα ECD and γc ECD, in this valency-sensitive format, with IC_50_s of 10 and 0.63 nM, respectively (Figs 2A and 2B). The IC_50_s of the peptide monomers, heterodimer, and MDK-703 bind to cynomolgus receptor subunits with affinities within 2-fold those of the respective human subunits. Direct binding affinities of MDK1472 for human IL-7Rα and γc subunits were determined by biolayer interferometry (BLI), yielding K_D_s estimated to be 302 and 16.5 nM respectively, consistent with the competition ELISA values (Figs 2C and 2D).

**Fig 2.**
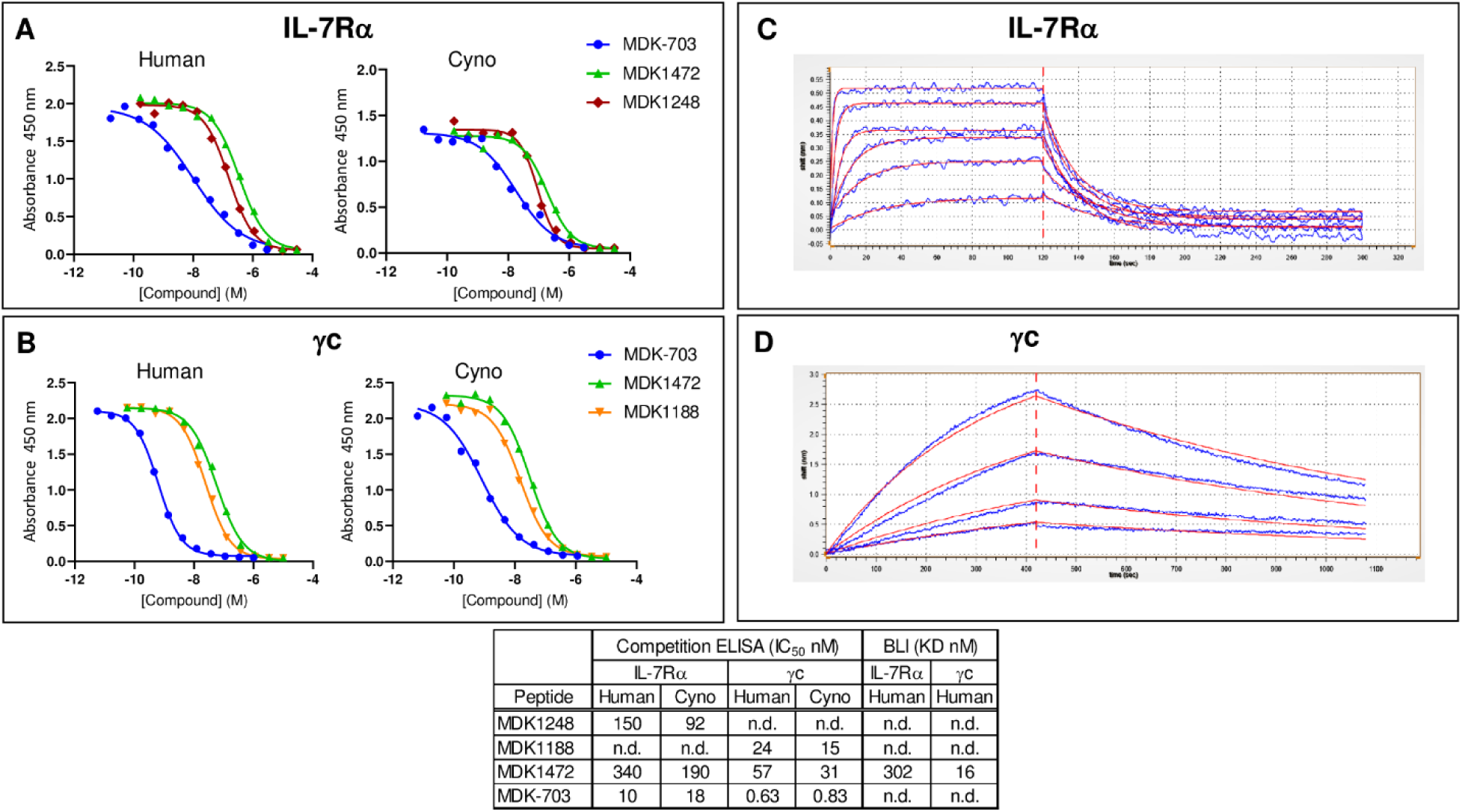
Binding of MDK1472, MDK-703, and component peptides to human and cynomolgus IL-7R subunits. (A, B) Determination of IC_50_ values in a competition ELISA for compound binding to human and cyno IL-7Rα and γc. The assay format is IL-7Rα-(His)_6_-tagged ECD or Fc-γc ECD immobilized on 96-well plates. Tracers are C-terminal biotinylated forms of the reference IL-7Rα and γc peptide ligands, each pre-complexed with Neutravidin-HRP (NA-HRP). (C, D) Label-free measurement of the equilibrium dissociation constant (K_D_) for peptide binding to IL-7Rα and γc extracellular domains with peptides was performed by biolayer interferometry as detailed in methods. Red lines depict 1:1 binding fit from Gator™ software.

Blue lines represent individual BLI sensogram data taken every 0.1 seconds. Inset table displays comparison of ELISA data and BLI kinetic data.

#### MDK1472 and MDK-703 act directly at the receptor, binding both subunits to induce IL-7R signaling

The synthetic peptide MDK1472 was designed to bind simultaneously to IL-7Rα and γc subunits and was tested for its ability to induce phosphorylation of STAT5, the primary IL-7R signaling pathway. An IL-7 responsive cell line, TF1-7Rα was constructed to express full-length human IL-7Rα in TF-1 cells, which naturally express γc, but not IL-7Rα. In a STAT5 phosphorylation assay, MDK1472 exhibits a potency of 5 nM and full efficacy (E_max_ equivalent to IL-7) (Fig 3).

**Fig 3.**
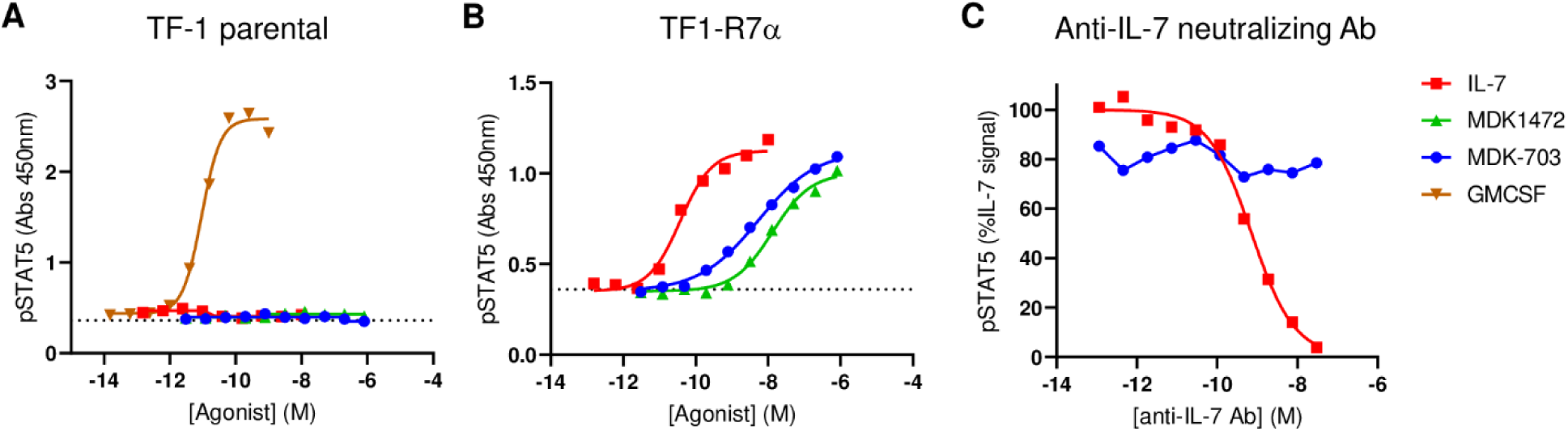
IL-7Rα dependence of agonist activity induced by MDK1472 and MDK-703 in TF-1 and TF1-7Rα cells: Compound-induced STAT5 phosphorylation in (A) TF-1 (γc+) and (B) TF1-7Rα (IL-7Rαγc+) cells. (C) pSTAT5 accumulation in TF1-7Rα cells treated with IL-7 or MDK-703 in the presence of an IL-7 neutralizing antibody. Cells were pre-incubated for 15 min with 1 pM to 30 nM anti-IL-7, and test compounds were then added at concentrations equivalent to EC_75_ and incubated for an additional 30 min. Reactions were stopped and cell lysates prepared for pSTAT5 analysis.

To unambiguously determine that MDK1472 acts by direct stimulation of IL-7R, we compared peptide-dependent receptor activation in the TF-1 parental and TF1-7Rα cells. As shown in Fig 3, agonist activity of IL-7, MDK1472, and MDK-703 are completely dependent on IL-7Rα expression. In addition, we repeated the pSTAT5 assays with TF1-7Rα cells in the presence of an IL-7 neutralizing antibody, showing that an antibody concentration that completely inhibits IL-7 stimulation has no effect on the activity of MDK-703, ruling out the possibility that IL-7R is activated by peptide-induced production of IL-7 by the cells.

### Receptor selectivity of MDK1472 and MDK-703

The cytokines and receptors of the common gamma (γc) chain family play essential roles in development, homeostasis, and regulation of immunity. Each family member uniquely contributes to shaping the immune response, and because each of the six receptors of the family depends on the availability of γc for normal signaling, it is important to understand any effects of MDK1472 and MDK-703 on the biology of the γc receptor system [38, 39]. In addition, a cytokine that is not a member of the γc family, thymic stromal lymphopoietin (TSLP) utilizes the IL-7Rα subunit as a co-receptor with CLRF2 (TSLPR), the private subunit of the TSLP receptor. TSLP exhibits a broad range of functions, targeting both lymphoid and myeloid cells, including CD4+ and CD8+ T cells, NKT, innate lymphoid 2, B cells, dendritic cells, neutrophils, mast cells, eosinophils [40]. Because TSLP activity depends on interaction with IL-7Rα, we examined the effect of MDK-703 on TSLP receptor-mediated events. We evaluated the potential for interference by MDK1472 and MDK-703 with the signaling of cytokine receptors utilizing γc or IL-7Rα subunits.

#### MDK1472 and MDK-703 bind both subunits in the IL-7R activation complex at sites different from IL-7

To gather information on the location of the MDK1472/-703 binding sites on the IL-7R subunits, we determined whether the novel agonists compete with natural IL-7 in activating the receptor. Because MDK1472 and MDK-703 are themselves agonists of IL-7R, we tested the individual (non-agonist) subunit binding domains of MDK1472, peptides MDK1248 and MDK1188 (Fig 1A), for inhibition of IL-7R-mediated receptor activation.

Fig 4A displays the dose-response of STAT5 activation induced by IL-7, MDK1472, and MDK-703 in the IL-7 responsive cell line TF1-7Rα. This assay shows IL-7, MDK1472, and MDK-703 to exhibit comparably efficacious pSTAT5 responses, with potencies ranging from 60 pM to 12 nM, as shown (note the individual peptide ligands MDK1248 and MDK1188 exhibit no induction of pSTAT5 at 10 μM). We next treated the cells with IL-7 (at ∼EC_75_) in the presence of saturating concentrations of MDK1248 or MDK1188, showing neither peptide to inhibit IL-7-induced pSTAT5 accumulation, indicating that these two peptides do not compete with IL-7 for binding to either receptor subunit, and that the interaction of MDK1472 (and MDK-703) with each subunit of IL-7R must occur at sites independent of those bound by IL-7 (Fig 4B).

**Fig 4.**
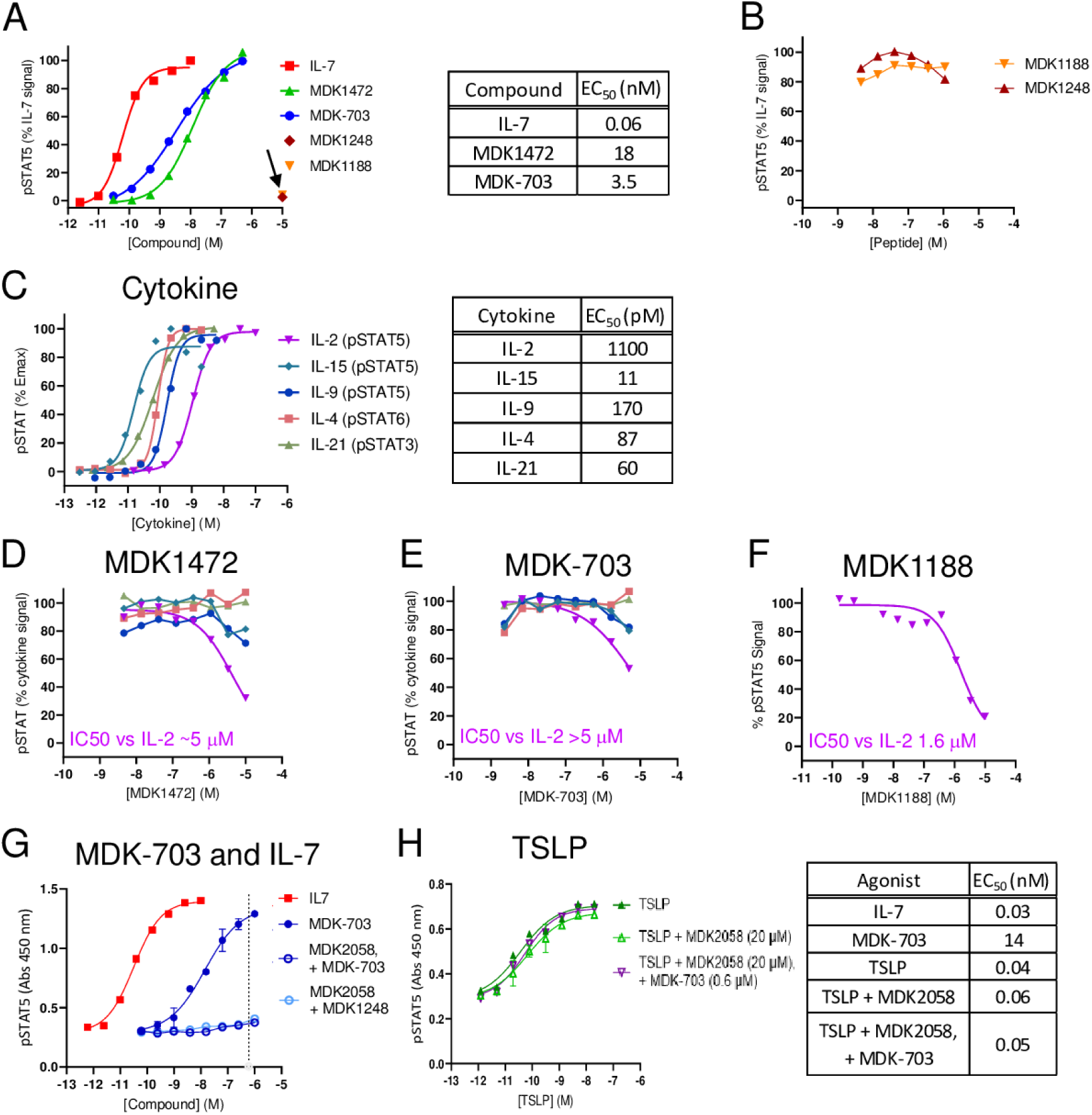
Effects of MDK1472, MDK-703, and their subdomain peptides on activation of receptors sharing subunits with IL-7R. (A) pSTAT5 dose response of IL-7 and the test compounds in TF1-7Rα cells (10 µM MDK1188 or MDK1248 alone indicated by arrow). (B) Response to IL-7 in the presence of 10 μM of each of the subunit-binding peptides MDK1248 and MDK1188 (the peptide fragments of MDK1472, which bind to IL-7Rα and γc, respectively). (C) Dose-response of IL-2, IL-4, IL-9, IL-15, and IL-21-induced STAT activation in TF-1 cells expressing the respective α or β receptor subunits. Inhibition by (D) MDK1472, (E) MDK-703, of the activation of each member of the γc receptor family by its respective cytokine agonist, and (F) IC_50_ of MDK1188 on IL-2. (G) Dose responses of IL-7 and MDK-703 induction of pSTAT5 in TF-1-7Rα/TSLPR cells, and MDK-703 in the presence or absence of MDK2058, a high affinity (IC_50_ = 300pM) competitive inhibitor of MDK-703 binding to the γc subunit. (H) Dose response of TSLP induction of pSTAT5 in TF-1-7Rα/TSLPR cells in the presence or absence of 20 µM MDK2058 and 600 nM MDK-703.

#### Effects of MDK1472 and MDK-703 on natural cytokine activation of γc family receptors other than IL-7R

The next series of experiments addressed the potential of MDK1472 and MDK-703 to interfere with γc cytokine receptors of IL-2, IL-4, IL-9, IL-15, or IL-21. We constructed a set of TF-1-derived cell lines, each responsive to a γc family cytokine: IL-4 (TF-1 naturally expresses both γc and IL-4Rα), IL-2 and IL-15 (engineered expression of IL-2/15Rβ), IL-9 (engineered expression of IL-9Rα), and IL-21 (engineered expression of IL-21Rα). Fig 4C displays the potency and efficacy of phosphorylation of the appropriate STAT in each cell line upon cytokine induction in the absence or presence of MDK1472 or MDK-703.

To assess any antagonistic effects of the test compounds, we treated each cell line with up to 10 μM MDK1472 or 5 μM MDK-703 in the presence of a sub-maximal level (∼EC_75_) of each cytokine. Only in the case of IL-2 stimulation of TF-1 cells expressing exogenous IL-2/15Rβ did we detect any interference by the test compounds: inhibition of pSTAT5 response with an IC_50_ of >5 μM (Figs 4D, E). We further determined that the IL-2R inhibition was produced by the γc ligand peptide MDK1188 alone (Fig 4F), establishing that activation of IL-2βγc receptor is weakly affected by MDK1472 and MDK-703 binding to γc subunit. The potency of this inhibition is ∼100-fold weaker than the affinity of MDK1472 (and MDK1188) for γc binding, indicating that inhibition of IL-2R activation by the test compounds is not competitive with IL-2, suggesting an allosteric effect on IL-2 receptor activation. Because the agonist potency of MDK1472 and MDK-703 on IL-7R is 3 to 4 logs greater than its IC_50_ for IL-2R inhibition, there should be no practical effect on the IL-2R system from therapeutic uses of MDK-703 as an IL-7R agonist (see below). It is notable that neither MDK1472 nor MDK-703 exhibit inhibition of IL-15, which shares the IL-2/15Rβγc receptor with IL-2. There is no detectable effect of the novel agonists on the activation of any of the other receptors of the γc family.

#### MDK-703 does not induce nor inhibit TSLP receptor activation

To assess the effect of MDK-703 on activity of the TSLP receptor, we prepared a TF-1-derived cell line engineered to express both IL-7Rα and TSLPR. These cells, designated TF-1-7Rα/TSLPR, are responsive to both IL-7 and TSLP in inducing STAT5 activation. As shown above, MDK-703 binds to IL-7Rα with an affinity of 10 nM at a site not competitive with IL-7. To test the effect of MDK-703 on TSLP activation of the TSLPR/IL-7Rα heterodimeric receptor, a pSTAT5 dose response of TSLP in the presence of 600 nM MDK-703 (40 to 100-fold the EC50 of IL-7R activation by MDK-703 in these cells) was measured. To suppress activation of the IL-7 receptor in this assay, we included MDK2058, a high affinity inhibitor of the interaction of MDK-703 with the γc subunit, allowing us to isolate the effect of MDK-703 on the TSLP receptor. Fig 4G displays the dose response of MDK-703 ± inhibitor MDK2058, showing that IL-7R activation by up to 1µM MDK-703 is fully suppressed by 20 µM of the inhibitor.

Furthermore, the dose response of TSLP activation is not detectably affected by the presence of the γc inhibitor, MDK2058, nor by the addition of 600 nM of MDK-703. These results indicate that MDK-703 does not interfere with activation of the TSLP receptor, and binds IL-7Rα at a site independent of TSLP.

### Response of human lymphocytes to test compounds and IL-7

#### Comparison of the kinetics and efficacy of pSTAT5 and proliferation induced by IL-7 and the novel IL-7R agonists in human lymphocytes *in vitro*

The most prominent lymphocyte signaling pathway induced by IL-7 is the JAK-initiated cascade leading to STAT5 activation and consequent regulation of downstream transcription. We analyzed STAT5 activation in isolated naive CD4+ and CD8+ T-cells to compare signaling mediated by IL-7 and the test compounds MDK1472 and MDK-703. We first assessed induction kinetics over a time course of compound exposure. Saturating concentrations of each of the compounds, previously determined to elicit maximum response of pSTAT5 signaling, were applied to the cells and pSTAT5 levels monitored for up to 2 hours. The resulting time-response profiles served to compare the kinetics of signal induction for the different compounds, and to determine T_max_ for correct time of exposure in subsequent dose response determinations for each compound. In these studies, IL-7-induced pSTAT5 accumulation rose rapidly to a maximum by 15 to 30 minutes, then declined slowly for the remainder of the time course. The rate of induction and T_max_ were found to be the same for IL-7, MDK1472, and MDK-703 in both cell types (Fig 5A). Dose response assays, conducted at 15 minutes of compound exposure (Tmax), revealed that MDK1472 and MDK-703 behave as full agonists in both cell types (Fig 5B), indicating that under the conditions of these studies, STAT5 pathway activation is similar in both rate of induction and efficacy upon treatment with IL-7 and the test compounds.

**Fig 5.**
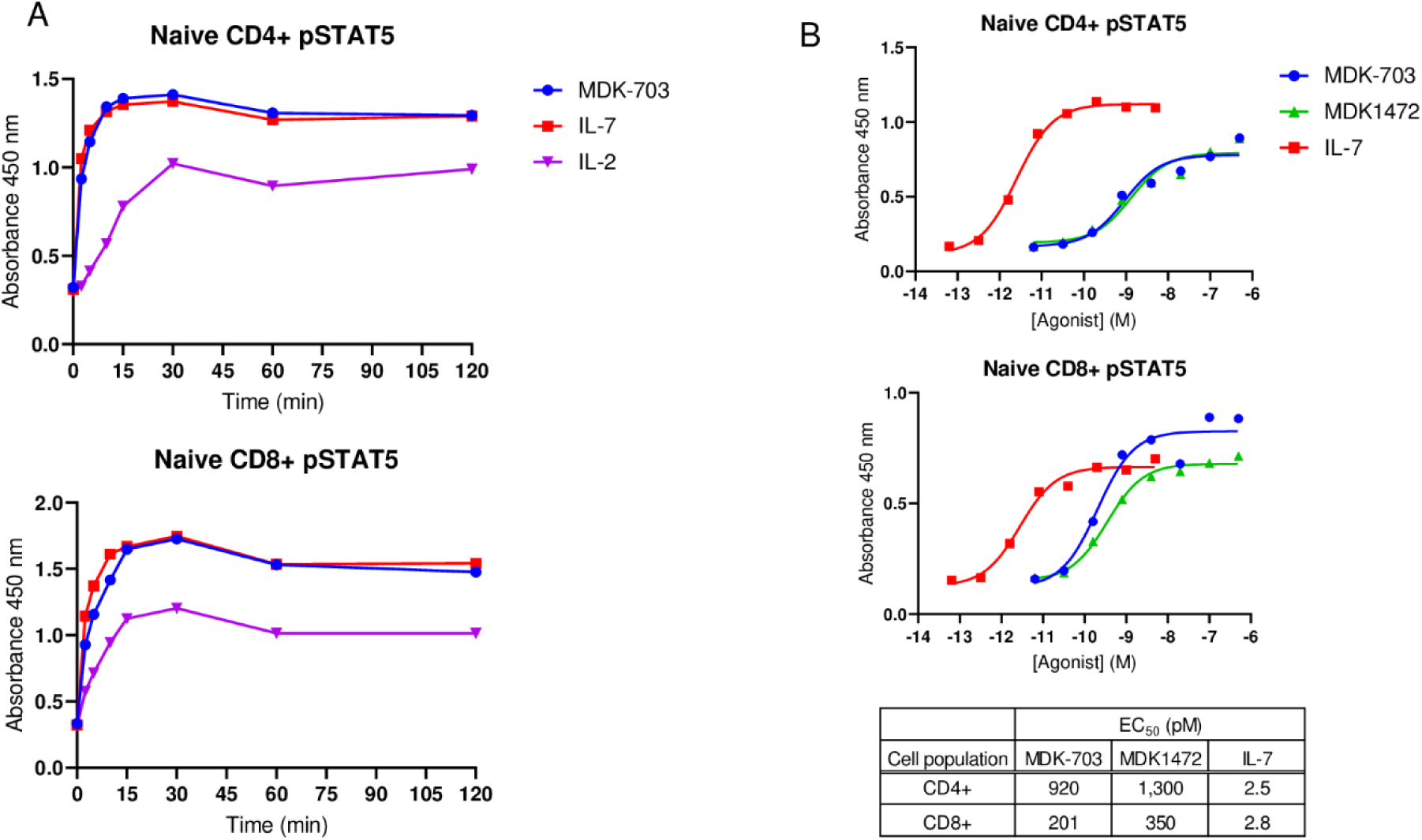
MDK1472 and MDK-703 behave as full agonists in activating the JAK-STAT5 pathway. Naive CD4+ and CD8+ T-cells, were separately isolated by negative selection from PBMCs, and rested overnight prior to compound exposure. (A) Cells were treated with saturating concentrations of IL-7 (10 nM), MDK1472 (1 μM), and MDK-703 (1 μM) for up to 2 hours and scored for pSTAT5 accumulation by ELISA. (B) The dose response for each compound was measured in human naive CD4+ and CD8+ cells, utilizing an exposure time of 15 min (T_max_).

#### MDK-703 exhibits cell selectivity similar to IL-7 in stimulating signaling and proliferation in major human T-cell populations

IL-7 exerts a broad range of effects on the lymphoid system, as most mature T-cells exhibit some IL-7Rα expression, with surface levels of IL-7Rα subunits and biological response to IL-7 fluctuating with the environment and differentiation status of the cells. To compare cell selectivity of MDK-703 with IL-7, we followed compound-mediated phosphorylation of STAT5 in major peripheral T-lymphocyte subpopulations of rested and CD3/CD28-activated PBMCs from 5 healthy donors following exposure to the compounds. Fig 6 displays dose-response curves for both compounds, assessed by flow cytometry of pSTAT5 gated on CD8+ and CD4+ T-cells, CD4+ regulatory T-cells (Treg), and NK cells. In each population, when rested or activated, the potency of IL-7 is significantly higher, but the efficacy (E_max_) of MDK-703 or IL-7 is the same, demonstrating that MDK-703 behaves as a full agonist of pSTAT5 activation in lymphocytes (Figs 6A and 6B). Consistent with the known functions of IL-7 agonists, both compounds induce the most robust pSTAT5 induction in CD4+ and CD8+ cells, with a significantly lower response in Tregs, cells that typically express low levels of IL-7Rα [41, 42]. Also as expected, NK cells show no pSTAT5 response to either IL-7 or MDK-703.

**Fig 6.**
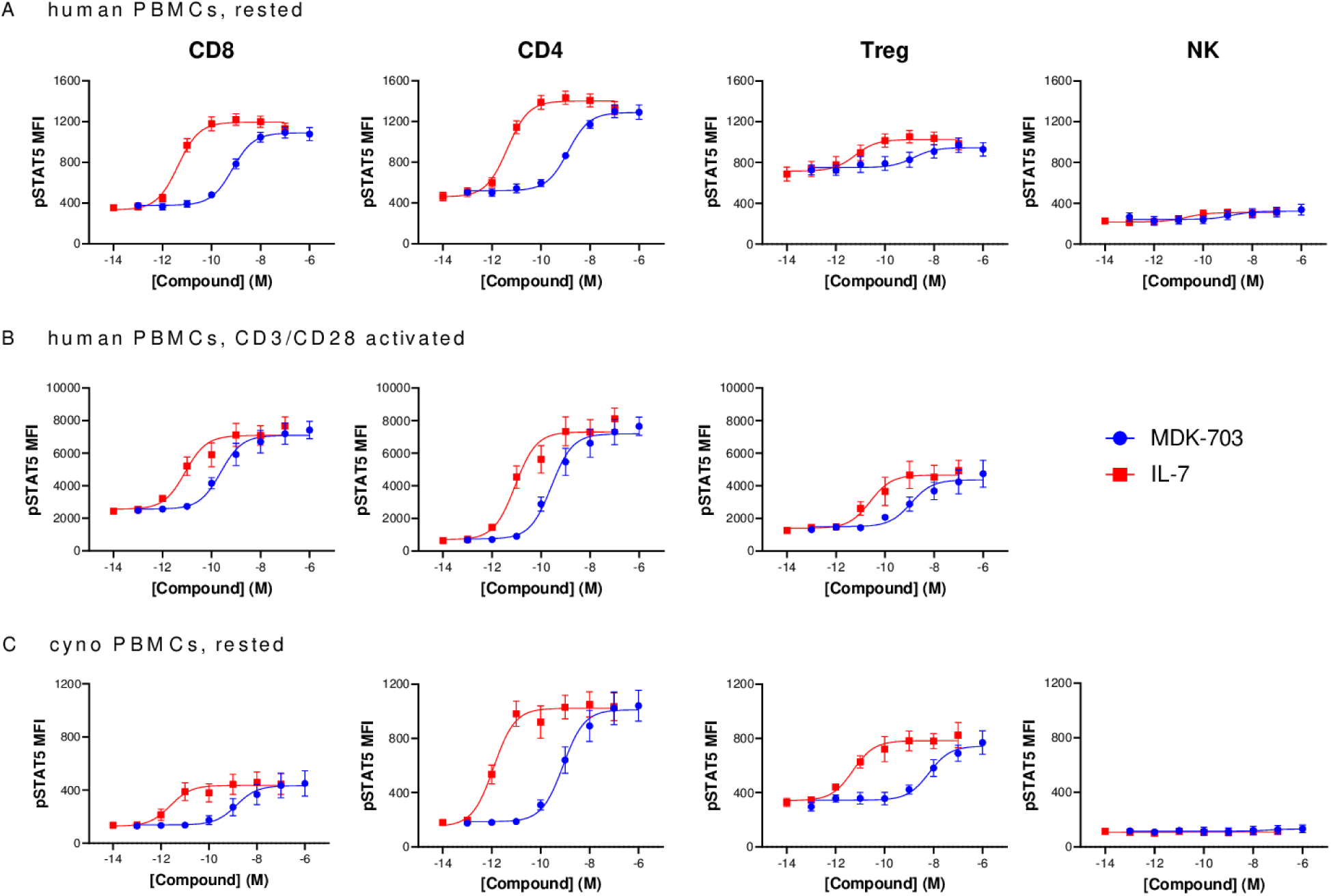
Induction of pSTAT5 in PBMCs treated with MDK-703 or IL-7. Frozen PBMCs from 5 healthy human donors (A, B) and cynomolgus monkey (C) were rested (A, C) or activated with anti-CD3/CD28 (B). Cells were stained for viability, followed by cell surface antibody staining on ice. Cells were washed and incubated with MDK-703 or IL-7 for 30 min at 37°C to activate the IL-7 receptor. After washing, cells were fixed, permeabilized, stained with anti-pSTAT5 antibody, and analyzed immediately by flow cytometry. The mean fluorescence intensity (MFI) of pSTAT5 detection was shown as mean ± SEM. (D), EC_50_ values of MDK-703 vs. IL-7 in immune subsets in human and cynomolgus PBMCs are shown in S1 Table. Flow cytometry gating data shown in S2 Fig.

We also evaluated pSTAT5 signaling in PBMCs of cynomolgus macaques, with results similar to the human studies: MDK-703 activated IL-7 receptor signaling in CD8+ and CD4+ T-cells, less in Tregs, and showed no activation of NK cells from the rested cynomolgus monkey PBMCs. Each compound exhibits potency comparable to that seen in human lymphocytes. As was the case with human PBMCs, both MDK-703 and IL-7 exhibit equivalent efficacy in cynomolgus lymphocyte subtypes (Fig 6C). These results demonstrate that MDK-703 is a full agonist of IL-7R with cell selectivity similar to that of IL-7.

An essential feature of IL-7-mediated homeostatic lymphoid maintenance is the proliferation of naive T-cells, expanding both recent thymic emigrants and circulating T_N_ clones to sustain the population and preserve a diverse TCR repertoire in the naive T-cell compartment. Memory T-cells require periodic IL-7 stimulation for long-term maintenance of the population and the potential for rapid expansion in response to antigen re-stimulation [43-45]. We treated rested PBMCs with MDK-703 or IL-7 (at E_max_ based on pSTAT5 dose-response) and determined Ki-67 levels, a marker of proliferative potential, and cell numbers of the major lymphocyte populations by flow cytometry (Fig 7). We found Ki-67 induction in each of the scored cell subsets to be similar upon treatment with MDK-703 or IL-7. In the CD8+ population, Ki-67+ frequency reached a peak by day 7, declining slightly by day 16. CD4+ T-cells exhibited a somewhat lesser response which was maintained to day 16. CD4+ Tregs, which typically express less IL-7Rα and are less sensitive to IL-7, displayed a significant rise in Ki-67 induction, but over a strong background of Ki-67 increase in the untreated cells. NK cells also exhibited a response profile similar to that of the CD4+ T-cell population; and as with Tregs, NK exhibited high Ki-67 background in untreated cells. The high Ki-67 background in untreated samples of Treg and NK cells may indicate a high basal turnover rate of these cell types in culture [46].

**Fig 7.**
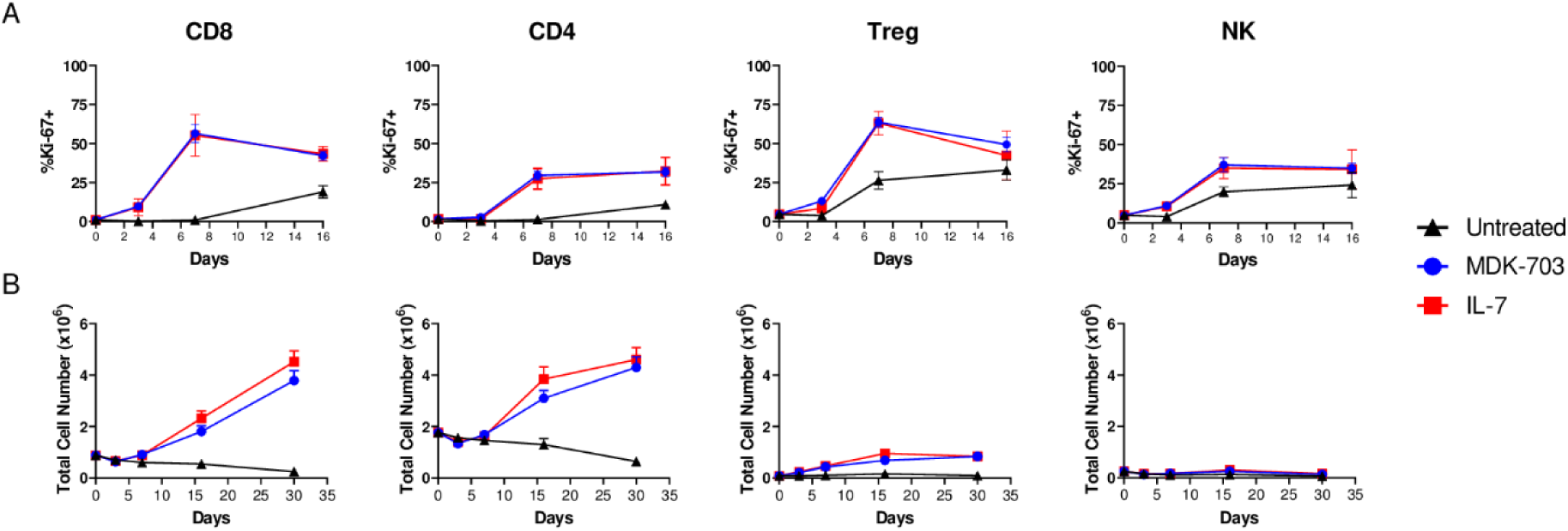
Proliferation of PBMC subpopulations following treatment with MDK-703 or IL-7. Frozen PBMCs from 5 healthy donors were rested overnight and treated with 100 nM MDK-703 or 1nM IL-7, or no added compound, and cultured for 30 days in the presence of compounds. On days 3, 7, 16, and 30, cell aliquots were taken and analyzed by flow cytometry for (A) Ki-67 expression and (B) absolute numbers of CD8^+^, CD4^+^, Treg, and NK cells. Data are shown as mean ± SEM. Flow cytometry gating data shown in S2 Fig.

In following treatment-dependent expansion of CD8+ and CD4+ T populations, we found CD8+ T-cell numbers were not increased by day 7 but exhibited strong expansion between days 7 and 16 of compound exposure and continued to increase through day 30 in culture, resulting in 4.3-fold (MDK-703) and 5.2-fold (IL-7) increases over baseline. The CD4+ T-cell population also expanded with similar kinetics, but with lower fold-increase over baseline by 30 days of compound exposure.

In the untreated samples, both CD8+ and CD4+ T-cell numbers declined over time. In the presence of the compounds, a slight increase in Treg cell numbers was observed; and NK cells showed no evidence of population expansion in this experiment, a result consistent with the known low response of NK cells to IL-7R agonists.

#### MDK1472 and MDK-703 induce IL-7Rα internalization with kinetics similar to IL-7

Upon binding to its receptor, IL-7 induces both signaling, and internalization of the active receptor complex. CD8+ cells in the absence of IL-7 continuously internalize IL-7Rα at a low rate via endosomal uptake with recycling to the cell surface; However, in the presence of IL-7 clathrin-mediated uptake of the signaling complex increases substantially, with IL-7Rα directed to late endosomes and targeted for proteasomal degradation [47, 48]. This reduces IL-7Rα surface expression and lowers cell responsiveness to IL-7. Internalization of the ternary signaling complex also depletes the local milieu of IL-7, an effect described as the “consumption” model of IL-7-mediated T-cell homeostasis [10, 11].

Because of the importance of IL-7Rα internalization and IL-7 consumption in modulating T-cell homeostasis, we evaluated the kinetics of receptor internalization induced by MDK1472 and MDK-703 in rested human CD8+ cells compared with IL-7 treatment. In previous studies, the decline of IL-7Rα on the surface of IL-7-treated CD8+ cells was reported to proceed with approximately linear kinetics, reaching ∼80% disappearance of receptor over 6 hours at 37°C [47]. We performed a similar experiment, tracking CD8+ surface IL-7Rα with a labeled anti-CD127 (IL-7Rα) antibody that we showed not to compete with either IL-7, MDK1472, or MDK-703 in binding and detecting the receptor subunit (S3 Fig). Surface IL-7R was pre-loaded with saturating concentrations of the agonists on ice and shifted to 37°C to allow internalization, then shifted back to ice to stop internalization. Negative controls consisted of untreated (no compound added); MDK1169, a peptide agonist of another γc family receptor (which contains a γc binding domain similar to MDK1472 but lacking the IL-7Rα binding domain); and MDK-202, an Fc-fusion of MDK1169 (a construct analogous to MDK-703, with the same Fc domain fusion partner). A parallel assay with all incubations held on ice for the duration of the time course was also run.

As shown in Fig 8 (and S3 Fig), IL-7 induces a temperature-dependent decline in IL-7Rα surface density, reaching a loss of 60% at 6 hours, consistent with the results of Faller, et *al.*[47]. The rate of surface decline of IL-7Rα upon treatment with MDK1472 or MDK-703 tracked very closely with that induced by IL-7. The negative control samples exhibited no IL-7Rα uptake at 37°C; and all samples held under cold conditions showed no change in IL-7Rα surface density. These observations are consistent with the interpretation that IL-7Rα disappearance from the cell surface upon treatment with these agonists is the result of cellular internalization, and the conclusion that MDK1472 and MDK-703 induce the uptake of IL-7Rα in rested CD8+ cells at a rate, and to an extent essentially identical to that driven by IL-7.

**Fig 8.**
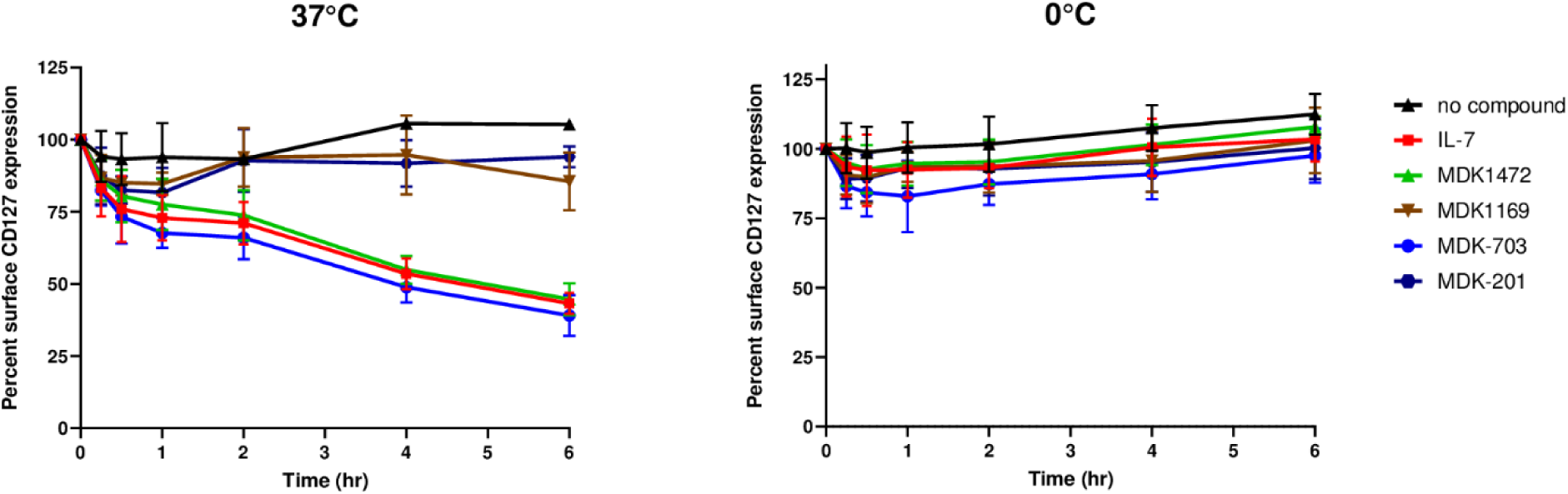
Test compounds induce decrease in cell surface IL-7Rα. Rested human CD8+ T-cells were treated with saturating concentrations of the test agonists (10 nM IL-7; 1 μM MDK1472 or MDK-703) or the negative control compounds (untreated, 1 μM MDK1169, or MDK-202) for 20 minutes on ice, then incubated at (A) 37°C or (B) 0°C for varying times to monitor uptake of IL-7Rα from the cell surface. Following timed incubations, the samples were stained, fixed, and analyzed by flow cytometry; and data was collected as median fluorescence intensity (MFI), as detailed in Methods. These data were normalized with the blank value (no added compound) set at 100% surface IL-7Rα, and the signal baseline set at 0. The primary flow data (MFI) is shown in S3 Fig.

### Effects of IL-7 and MDK-703 on T-memory subpopulations

#### MDK-703 drives expansion of naïve and memory T-cell subpopulations in human PBMCs

Naive and memory subsets of peripheral T-cells are highly responsive to IL-7 by virtue of robust expression of IL-7Rα. We compared the effect of IL-7R agonists IL-7 and MDK-703 on CD8+ T-memory subpopulations in rested human PBMCs, and found the effect of MDK-703 on cell number to be similar to that of IL-7 on T-naïve and on all Tmem populations observed (Fig 9).

**Fig 9.**
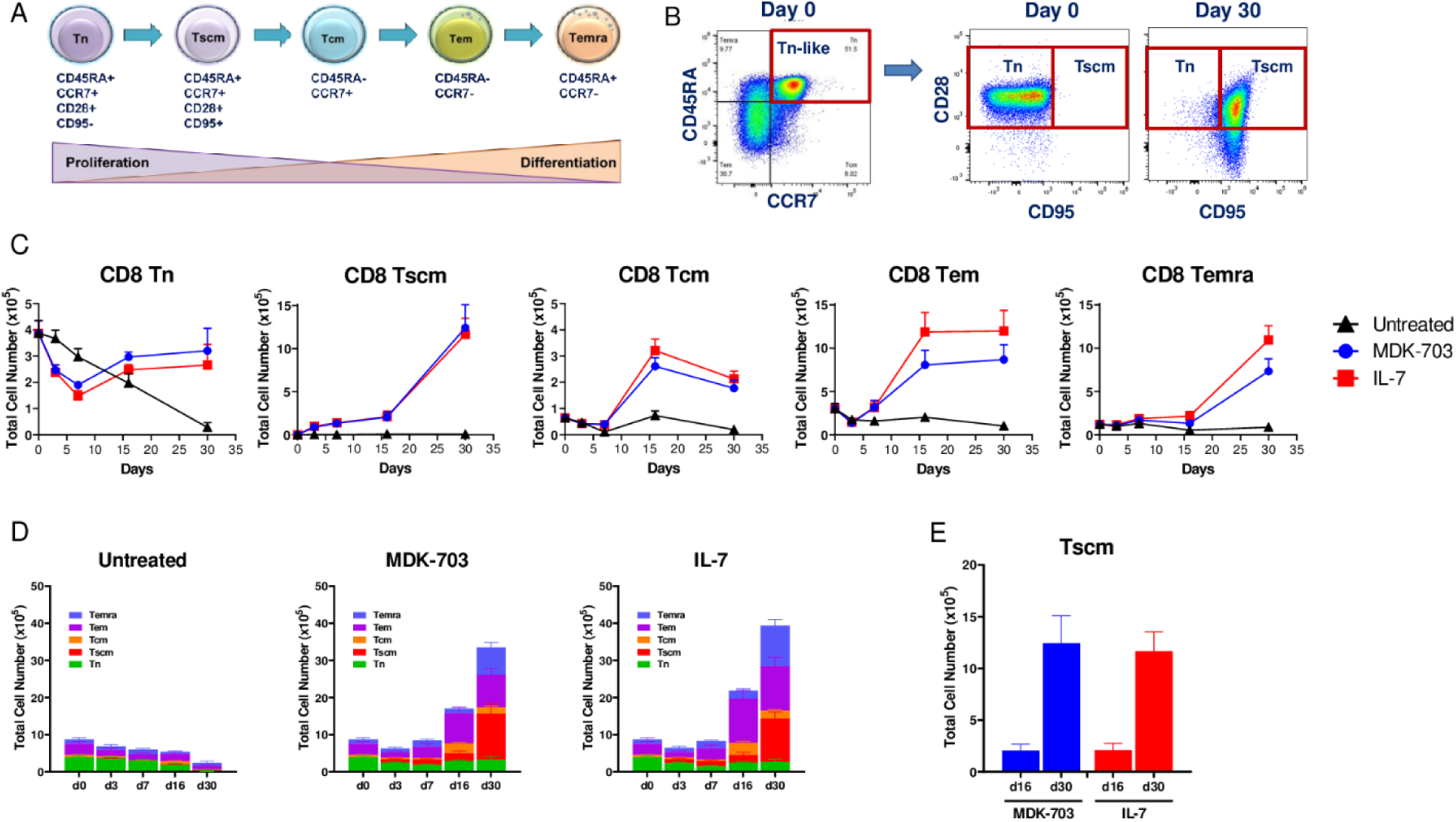
The expansion of CD8+ T memory subpopulations following treatment with MDK-703 or IL-7. Frozen PBMCs from 5 healthy donors were rested overnight and left untreated or treated with 100 nM MDK-703 or 1 nM IL-7 in culture. On days 3, 7, 16, and 30, cell aliquots were taken and analyzed by flow cytometry of CD8+ T naïve and memory populations. (A) Schematic of the putative differentiation pathway of the memory T cell compartment. (B) differential gating of naïve and early memory (Tscm) subpopulations. (C) Cell counts of treated and untreated memory subpopulations over time. (D) Stacked bar representation of changes in total memory T cell subpopulations over time. (E) Comparison of CD8+ Tscm expansion on day 16 and day 30 of culture with MDK-703 or IL-7. Data are shown as mean ± SEM. Detailed gating information is provided in S4 Fig.

Treatment with both compounds caused an immediate decline in CD8+ naive cells, reaching a nadir by day 7, followed by a return to initial levels by day 16 that were sustained to the end of the 30 day duration of the experiment. In contrast, memory T cell populations were low at T_0_ and exhibited little or no initial dip, followed by substantial increase in cell count by day 16 of exposure to the compounds. Most notable is the behavior of a very early effector/memory subset designated T-stem cell memory (Tscm), a cell type exhibiting persistent stem-like properties (Fig 9B). Both MDK-703 and IL-7 drive an increase in CD8+ Tscm cell number beginning day three, and a dramatic population expansion to reach a 1300-fold increase by day 30, compared to a 5-fold expansion in the untreated group. The remaining memory populations exhibited more modest (fold) increases in cell number following exposure to either IL-7 or MDK-703. CD8+ central memory (Tcm) exhibited a modest initial decline, then a rapid rise to a peak by day 16 of 5-to 6-fold over day 0, followed by a slight decline by day 30 to 3-to 4-fold elevation over day 0. Similarly, CD8+ effector memory (Tem) exposed to the IL-7R agonists exhibited a small early decline but increased 3-and 4-fold by day 16, a level which persisted until day 30. CD8+ terminally differentiated effector memory cells (Temra) responded to IL-7 and to MDK-703 with marked expansion of 7-and 11-fold over 30 days. For naïve and all memory populations, cells showed little evidence of expansion in the absence of IL-7R agonists (untreated) These results demonstrate MDK-703 to be as effective as IL-7 in stimulating memory T cell expansion, in particular the stem-like Tscm population (Figs 9B and 9C). The profound effect of both IL-7 and MDK-703 on the cell number of the overall memory compartment as well as the individual subpopulations is illustrated in the histograms shown in Figs 9D and 9E.

#### MDK-703 drives expansion of immune cells in huCD34^+^-engrafted NSG mice

MDK-703 binds and activates both human and NHP IL-7 receptors but is inactive on the murine receptor. To evaluate the effects in a rodent model, we utilized NSG mice engrafted with human CD34+ hematopoietic stem cells (CD34+ HIS mice) (Fig 10). Mice were administered a single intravenous dose of human Fc isotype control or MDK-703 at 1 mg/kg. Peripheral blood samples were analyzed on day 7, and terminal blood and splenocytes were analyzed on day 12 .

**Fig 10.**
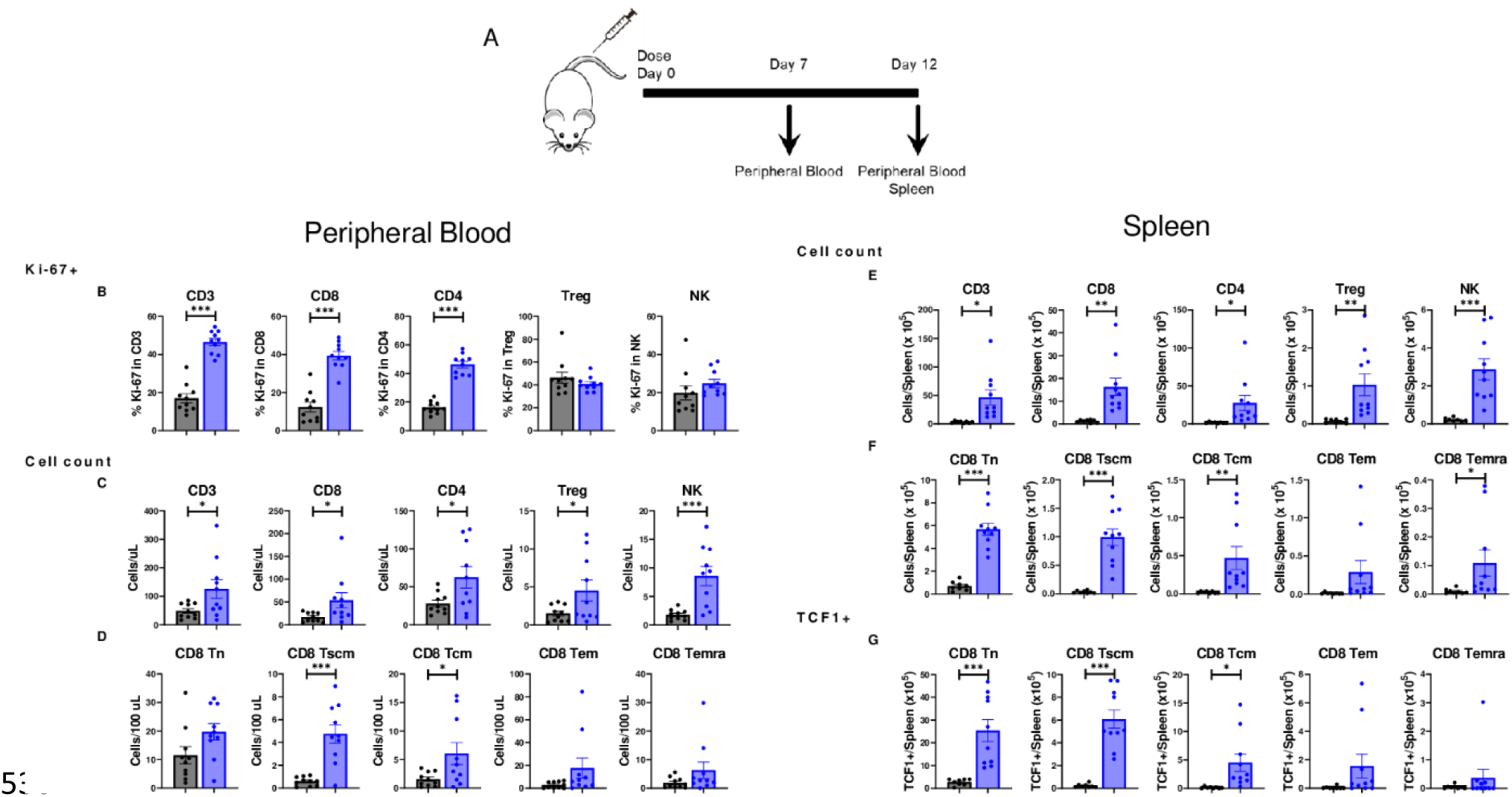
Expansion of immune cell subpopulations in humanized mice treated with MDK-703. NSG mice (n=10 per treatment) engrafted with human CD34+ cells from two donors (5 mice/donor) were dosed once intravenously with 1 mg/kg Fc (gray bars) or MDK-703 (blue bars), and peripheral blood (B-D) and spleen (E-G) were collected and analyzed at the indicated times by flow cytometry. (A) Diagram of the experimental plan. (B) Frequencies of Ki-67+ in CD3+, CD4+, CD8+, Treg, and NK cell populations in peripheral blood on day 7. (C) Absolute cell numbers of CD3+, CD4+, CD8+, Treg, and NK cell populations in peripheral blood on day 12. (D) CD8+ T memory subpopulations in peripheral blood on day 12. (E) Absolute cell numbers of CD3+, CD4+, CD8+, Treg, and NK cell populations in the spleen on day 12. (F) CD8+ T memory subpopulations in the spleen on day 12. (G) TCF1 expression in CD8+ T memory subpopulations in the spleen on day 12. Population gates were drawn based on FMO controls. Statistical analysis was done using Student’s T-Test. *p<0.05, **p<0.005, and ***p<0.0005. Detailed gating information is provided in S6 Fig.

Consistent with the pSTAT5 induction data, MDK-703 increased the Ki-67+ population in peripheral CD8+ and CD4+ T-cells, but not in Tregs or NK cells on day 7 (Fig 10B); however, at this time point, cell numbers were not increased over the Fc control group (S5 Fig). On day 12, Ki-67+ cells were no longer observed (S5 Fig), but CD8+ and CD4+ T-cell numbers were elevated (Fig 10C). In vivo, both Treg and NK cell numbers, typically unresponsive to IL-7 agonists, were increased, suggesting an indirect effect of MDK-703, possibly driven by IL-7R-dependent release of cytokines in the engrafted immune system.

The CD8+ Tn and Tcm compartments showed elevated frequencies of Ki-67+ cells on day 7 (S5 Fig), and Tn, Tscm, and Tcm cell numbers expanded in response to MDK-703; but Tem and Temra did not reach significance – an observation possibly due to the limited time course of this experiment, as we had observed a robust in vitro expansion of Tem and Temra from PBMC in 16 to 30 days of culture (see Fig 9).

More dramatic changes were observed in the spleen. Absolute cell numbers were significantly increased in CD8+, CD4+ T-cells, Treg, and NK cell populations (Fig 10E) without changes in the Ki-67+ except for CD4+ T-cells (S5 Fig) by MDK-703. The effect on NK cells is assumed to be a secondary or indirect effect of MDK-703 treatment. CD8+ naïve and memory subpopulations were expanded to a greater extent in the spleen than in blood by 12 days of treatment with MDK-703 (Fig 10F). We observed Tn, Tscm, Tcm, and Temra, to be significantly expanded (Tem did not reach significance). T-cell factor 1 (TCF1) transcription factor is expressed in self-renewing immune cells. Since TCF1+ expression is found in naïve and Tscm cells, we examined the effect of MDK-703 on TCF1+ cell expansion. As shown in Fig 10G, MDK-703 treatment increased TCF1 frequency in CD8+ Tn and Tscm, which are highly proliferative and less differentiated, less in Tcm; but non-significant in Tem and Temra, cells which are more differentiated and have lower proliferation capacity. The results indicate that MDK-703, like IL-7, drives a substantial increase in stem-like memory cells.

### MDK-703 in non-human primates

#### MDK-703 exhibits an extended circulating half-life and induces persistent elevation of peripheral lymphocytes in non-human primates

MDK-703, an Fc-fusion designed to extend the half-life of the pharmacologically active IL-7 agonist peptide MDK1472, was administered as a single dose of 1 mg/kg to naive cynomolgus macaques via intravenous, subcutaneous, or intramuscular injection. MDK-703 was well tolerated, with no adverse effects on clinical observations or blood chemistry.

The mean terminal half-life of MDK-703 was 46 hours in these animals, with higher bioavailability achieved after subcutaneous or intramuscular injection (Fig 11A). In a separate study, three animals were dosed once subcutaneously with 0.3 mg/kg and observed for changes in absolute lymphocyte count (ALC, hematology) and the expansion of CD8+, CD4+, Treg, and NK cells (by flow cytometry) for up to 42 days. Following administration, absolute lymphocyte (ALC) counts dropped from baseline to a nadir at 1-2 days, recovering to baseline by day four and reaching a peak of ∼130% over baseline by day 10 (Fig 11B). ALC remained elevated through day 29, and slowly declined to near baseline level by day 41, demonstrating that the MDK-703 pharmacodynamic effect persists well beyond the exposure to treatment. The initial drop in cell number was observed in CD8+ and CD4+ T-cells, peaking on day seven and remaining elevated (>50% over baseline) for 29 days (Fig 11C). There was no significant increase in Treg count, consistent with the data from CD34+ humanized mice. The initial drop in blood lymphocytes following IL-7R agonist administration is attributed to the induction of adhesion molecule expression directing the uptake of responding cells into tissues [49].

**Fig 11.**
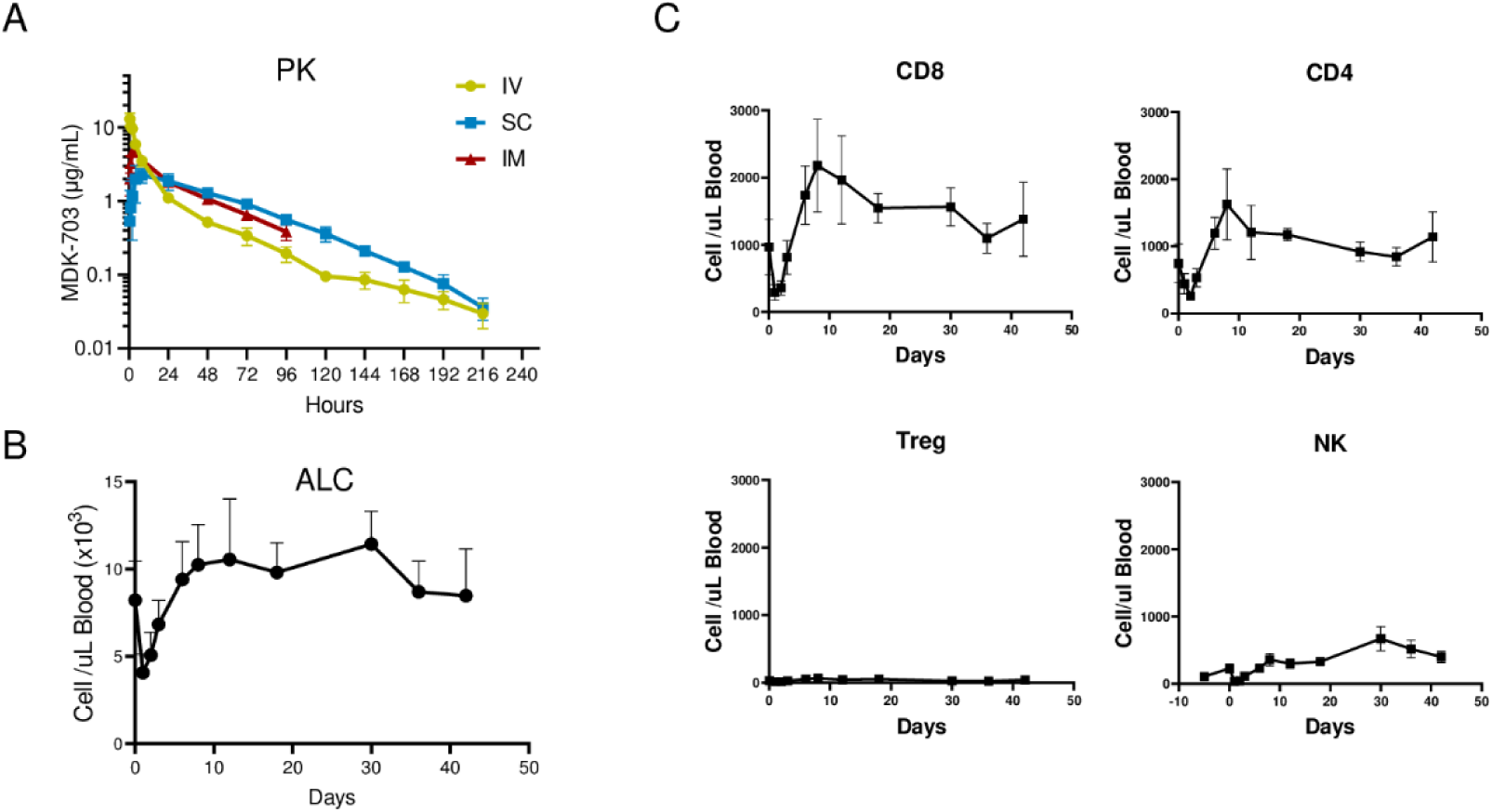
Pharmacokinetic (PK) and pharmacodynamic (PD) properties of MDK-703 in cynomolgus macaques. (A) PK of MDK-703: Animals (n=3) were administered a single dose of 1 mg/kg MDK-703 via IV, SC, or IM. The serum concentration of MDK-703 at the indicated time points was determined by sandwich ELISA. (B and C) PD effect of MDK-703: Three animals were dosed once subcutaneously with 0.3 mg/kg, and blood samples were collected at the indicated time points for absolute lymphocyte counts (B) and immune profiling of CD8, CD4, Treg, and NK cells by flow cytometry (C). Data show mean ±SEM.

Proliferation and expansion of lymphocytes then follow, with a duration of effect well beyond the circulating persistence of the compounds, consistent with the known protective, anti-apoptotic properties of IL-7R agonists in extending T-cell lifetime [50]. These results are similar to those reported in patients treated with a PK-stabilized formulation of IL-7 (GX-17; IL-7 fused to an Fc-domain) [25].

## Discussion

MDK-703 is the fusion of a structurally novel IL-7 peptidyl agonist with an huIgG2-Fc domain. The peptide component, MDK1472, was assembled as a single chain heterodimer of novel peptide ligands of IL-7Rα and γc joined by a short flexible amino acid linker. A notable feature of the active agonist peptide is the stochastic process of discovery of the component ligands, leading to an IL-7R agonist completely unrelated in sequence to native IL-7. We found MDK1472 to exert receptor agonism through binding to sites distinct from the binding sites of IL-7 on both receptor subunits. Although the relative affinities of MDK1472 for IL-7Rα and γc differ from that of IL-7, signal transduction initiated by the formation of a ternary complex with both receptor subunits is similar to that of IL-7.

The novel sequence of the active peptide provides distinct benefits over natural IL-7 as a therapeutic agent. Various forms of IL-7 modified for clinical applications exhibit immunogenic properties, including induction of antibodies which neutralize IL-7 activity [25]. Such IL-7 neutralizing antibodies have the potential to affect endogenous IL-7, which can produce significantly detrimental effects in patients. The small size of the peptide component of MDK-703 reduces the likelihood of harboring immunogenic epitopes, and *in silico* algorithms predict a very low propensity for immunogenicity. Furthermore, should an immunogenic response to the novel peptide sequence occur, ADAs would not neutralize endogenous IL-7. The single chain peptidyl structure of the peptide also provides a means of pharmacokinetic enhancement by fusion to an Fc partner; and the Fc -fusion MDK-703 exhibits substantial persistence in the blood of non-human primates after IV, IM, and SC administration.

Some studies on the clinical use of agonists derived from natural IL-7 report injection site reactions in patients. These effects may be explained by the reported binding of IL-7 to components of the extracellular matrix, potentially resulting in injection site depot effects [4, 51]. In addition, recombinantly expressed proteins occasionally form aggregates, as exemplified in the report of a fusion of IL-7 to IgG-Fc [52]. Although formulations can be employed which limit this effect, subsequent injection of high concentrations of formulated drug into physiological aqueous environments may induce aggregation and local deposition. MDK-703 injected SC or IM into NHP, exhibited systemic bioavailability similar to that produced by IV dosing. Preliminary assessment of MDK-703 dosed IM in healthy human subjects found no injection site reactions, and systemic MDK-703 exposure consistent with that predicted from NHP studies, suggesting high bioavailability in humans [53].

Because the peptidyl agonist component of MDK-703 is not naturally occurring, is unrelated to IL-7, and acts through a unique binding complex with IL-7R, we undertook a detailed comparison of receptor signaling and cellular responses induced by the synthetic peptide, the Fc-fusion, and IL-7. We showed MDK1472 and MDK-703 to activate the IL-7R through direct receptor interaction to faithfully emulate the kinetics and efficacy of IL-7 internalization, signal transduction, and stimulation of proliferation in human lymphocyte subpopulations. At concentrations that produce maximal IL-7R-mediated pSTAT5 elevation, neither MDK1472 nor MDK-703 significantly affected the activity of any other γc cytokines at their respective receptors.

To compare the biology of MDK-703 with IL-7, we measured pSTAT5 in the major lymphocyte subtypes CD4+, CD8+, CD4+Tregs, and NK cells in human and cynomolgus PBMCs; and found that the kinetics and Emax of pSTAT5 induction in each subtype is essentially the same for both MDK-703 and IL-7, and this finding held for PBMCs of both species. CD4+ and CD8+ cells exhibit the most robust response to the compounds, followed by more modest effects on CD4+Tregs, and very little response in the NK population --results consistent with previously described properties of IL-7R agonists. With respect to Tregs, IL-7R agonists have been reported to reduce their immunosuppressive properties [41, 54], potentially enhancing the therapeutic utility of MDK-703 in oncology and infectious disease.

The *in vitro* potency of MDK-703 in the pSTAT5 assay is 10 to 100-fold lower than that of IL-7. In some cases, attenuation of the potency of cytokine derivatives has been shown to increase the availability of drugs that are subject to target receptor-mediated disposition (TMDD) [55]; and we previously reported that the potency of MDK-703 is sufficient to achieve robust PD effects on blood T-cells at doses as low as 30 µg/kg IM in healthy human subjects [53].

In addition to *in vitro* pSTAT5 phosphorylation, we showed both MDK-703 and IL-7 to induce similar expansion of lymphocyte numbers in culture: CD4+ and CD8+ cells reached maximal counts 5-fold over day 1 baseline levels between days 7 and 16, levels which persisted for at least 30 days.

The effects of IL-7R agonists on naïve and memory T-cell compartments are well known and are critical for the central role of IL-7 in maintaining a healthy immune response. Exploring compound-dependent effects on these lymphocytes, we compared MDK-703 with IL-7 on rested human PBMCs and found the proliferative effects on each scored subpopulation to be very similar for both compounds. Focusing on CD8+ cells, we observed the number of naïve cells to decline quickly and substantially, reaching a nadir by day 7, followed by a return to day 0 basal level, which was sustained to day 30. In contrast, the memory T-cell populations (Tscm, Tcm, Tem, Temra) all exhibited expansion of their initial cell number, with the most prominent effect an early and dramatic increase of Tscm cells, a memory phenotype exhibiting evidence of both activation and persistent stem-like properties. Both compounds drove the immediate increase of Tscm number to reach an expansion of 1300-fold by day 30. More modest increases of Tcm and Tem numbers followed a slightly delayed time course, reaching 5 to 6-fold and 3 to 4 -fold over baseline, respectively, by day 16. These results demonstrate MDK-703 to be at least as effective in stimulating Tmem – particularly Tscm – expansion as IL-7; and serve as further evidence of the similarity in T-cell immuno-biology produced by these two IL-7R agonists *in vitro*.

MDK-703 exhibits substantial proteolytic stability in plasma *in vitro*, and persistence after administration to NHP with circulating T_1/2_ of 46 hrs. Administration by the IM and SC routes demonstrated an MDK-703 AUC similar to that of IV administration.

MDK-703 is active on primate IL-7 receptors but exhibits undetectable activity on rodent receptors. To study the pharmacodynamic properties on human immune cells, we utilized an *in vivo* model huCD34+ engrafted NSG HIS mice. A single IV dose of MDK-703 increased the frequency of Ki-67+ CD4+ and CD8+ T cells, but not Tregs nor NK cells, by day 7. On day 12, Ki-67+ cells were no longer present in the peripheral CD8+ and CD4+ populations, but the cell numbers were significantly increased. In these *in vivo* experiments, Tregs and NK cells, typically relatively unresponsive to IL-7R agonists, were modestly elevated. This expansion may be an indirect effect of L-7R-dependent release of cytokines other than IL-7, that stimulate these populations. Scoring for the Tn and Tmem subsets in blood showed CD8+ Tscm and Tcm cells to exhibit an increase in frequency of Ki-67+ expression at day 7, while Tn, Tem, and Temra did not. By day 12, the absolute cell numbers in Tn and all Tmem compartments were significantly increased in blood, and substantially elevated in spleen. The frequency of TCF1+ cells increased significantly in Tn, Tscm,Tcm, and Tem, These results demonstrate that MDK-703, like IL-7, drives a general increase in the number and “stemness” of memory cells.

In cynomolgus macaques, a single SC dose of 0.3 mg/kg produced an initial drop in blood lymphocytes, characteristic of the tissue trafficking effects of IL-7R agonists, followed by a return to baseline by day 4, and climbing to 230% of baseline by Day 11. This level remained elevated through day 29, and declined slowly thereafter to baseline at day 41, indicating persistence of the expanded lymphocytes well beyond the period of exposure to the compound.

Stimulation of the memory T-cell compartments appears to be of importance for the reported anti-tumor activities of IL-7R agonists, either administered to animals or patients; or treatment of engineered T-cells undergoing *ex vivo* expansion prior to transfer to patients. Amplification of Tscm, a naive-like, but antigen-experienced population that retains a minimally differentiated and highly proliferative self-renewal phenotype, appears to be a particularly important subset of the T-memory compartment in this regard. As a source of potent effectors resistant to antigen-driven exhaustion, cells of the Tscm subtype are predicted to exert a key influence on T-cell-mediated anti-tumor immunity. In animal tumor models, both CD4+ and CD8+ Tscm are implicated in anti-tumor efficacy; in humans, administration of CAR-T cells with enhanced Tscm phenotype leads to improved persistence of the engineered cells, as well as improved clinical outcome; and in healthy individuals Tscm exhibit greater proliferation and cytotoxic polyfunctionality with a higher frequency of the triple positive effector phenotype -- elevation of IFNγ, IL-2, and tumor necrosis factor (TNF)α secretion -- upon secondary antigen response, than do other Tmem-subsets [13-15, 27, 56-58].

MDK-703, the novel IL-7R agonist described here, produces strong stimulation of Tscm production in PBMC cultures, and rapid appearance in both blood and spleen of MDK-703 treated HIS mice. This property of MDK-703 appears to be translatable to humans, as a single dose of 30 µg/kg of MDK-703 in healthy subjects resulted in a 9-fold increase in CD8+ Tscm. [53]. The ability to increase the Tscm population may be an important attribute of the therapeutic utility of MDK-703.

In summary, MDK-703 exhibits properties that make it a strong candidate for clinical evaluation. These include its novel chemical composition to avoid IL-7 neutralizing immune response, high bioavailability, and extended half-life. Despite its novel composition and unique ternary complex with IL-7Rα and γc, MDK-703 appears to faithfully emulate the properties of IL-7 with respect to cellular signaling and *in vivo* expansion of T-cell subpopulations. Given the unique pharmacology of MDK-703, it may have utility in indications where its pharmacokinetics, lack of IL-7 neutralizing antibodies, and T-cell enhancing properties are beneficial, including oncology, anti-viral and other infectious disease, vaccine enhancement, and treatment of lymphopenia.

## Materials and Methods

### Ethics statements

#### Human primary tissue samples (PBMCs)

Buffy coats, PBMCs, and isolated PBMC subsets were obtained from the Stanford Blood Center (Stanford University, Stanford CA) collected under eProtocol “Minimal Risk Research Related Activities at Stanford Blood Center” (SQL 79075) #13942; IRB (registration 5236). Samples were collected with written donor consent and anonymized prior to provision for research purposes. Stanford University is in compliance with requirements for protection of human subjects, including 45 CFR 46, 21 CFR 50 and 56, and 38 CFR 16.

Human PBMCs were obtained from Stemcell Technologies (Vancouver, BC). Samples were collected from donors and anonymized prior to providing for research purposes. Stemcell Technologies provided Institutional Review Board (IRB)-approved consent forms and collection protocols for all lots of PBMCs used in these studies: “Leukapheresis Sample Collection from Healthy Human Donors for the Purpose of Scientific and Medical Research and education” (IRB protocols 2019110, 018-V.1; 018-V.7; 018.V.11; including written consent forms U5 and U9).

#### Human-derived established cell lines

Human-derived cell lines used in this study were initially obtained from accredited commercial suppliers: (1) HEK293 (human embryonic kidney cell line, FreeStyle® 293-F) was obtained from Invitrogen (San Diego, CA) for use as a recombinant protein expression vehicle; and (2) TF-1 erythroblast cell line was obtained from the American Type Culture Collection (ATCC Cat# CRL-2003) for use as the parental cell line for the engineered expression of each full length private receptor (alpha or beta) of the gamma common family, and CFLR2. No cell lines were prepared directly from human tissue at Medikine.

#### Mouse Studies

Mouse PK and PD studies were carried out at Crown Bioscience (San Diego, CA) in strict accordance the regulations of the Association for Assessment and Accreditation of Laboratory care (AAALAC). The study protocol was approved by the Committee on the Ethics of Animal Experiments of Crown Bioscience (Protocol number CDSD-ACUP-002). The subject animals (HuCD34+ humanized mice (NSG™ NOD.Cg-Prkdcscid Il2rgtm1Wjl, Stock No.005557) were provided directly to Crown BioScience by The Jackson Laboratory (Bar Harbor, ME).

#### Non-human primate studies

NHP (cynomolgus macaque) studies 2006 and 2102 were performed according to Primate Products, LLC (Miami, FL) Standard Operating Procedures, authorized veterinary standards and the present protocols. The protocols were approved by the Envol Biomedical IACUC (Naples, Fl) (Study 2006 Approval number 7117-21; Study 2102 approval number 7174-21). Prior to initiation, all animals underwent a physical examination by the study veterinarian. Upon study completion, the animals were placed in the stock colony of non-naïve animals held at Primate Products, LLC (PP LLC).

##### Housing, feeding, and environmental enrichment

Purina 5049 was provided daily in amounts appropriate for the size of the animal. Tap water was provided *ad libitum* via an automatic watering device. The animals were housed according to Primate Products, LLC (PP LLC) standard operating procedures. Each animal’s immediate holding cage was cleaned once daily. Enrichment was provided as needed.

##### Humane endpoints

No humane endpoints were included in this study. The study was designed to evaluate pharmacokinetics and pharmacodynamics. Animals were observed twice a day until the end of the study and returned to the stock colony.

#### Preparation of compounds: Peptides and Fc-fusions

##### Synthesis of peptides

All peptides (MDK1472, MDK1188, MDK1248, MDK1169, MDK2058) were synthesized using standard Fmoc/HATU/TFA synthesis protocols. For peptides containing two cysteines, the single disulfide bond was formed using trityl protection for the thiols and dimethyl sulfoxide (DMSO) oxidation. For peptides containing four cysteines, and with the potential of forming three disulfide bond arrangements, orthogonal protecting groups were used for the two pairs of cysteines to ensure formation of only the desired disulfide bridges: one cysteine pair protected with trityl and oxidized with DMSO, and the second pair protected with Acm and oxidized with iodine. Peptides were purified by preparative reverse phase HPLC (≥95%), and correct mass verified by ESI-MS.

#### Preparation of MDK-703

A mammalian expression vector was constructed to express peptides linked to Fc-fragments consisting of the CH2 and CH3 domains of the heavy chain and hinge regions of human IgG2 Fc modified as follows: The first and second cysteines of the hinge region were replaced with serine to prevent detrimental disulfide bridge formation; the last amino acid (lysine) of the Fc region was replaced with an alanine for fusion stability. This vector contains the strong constitutive cytomegalovirus (CMV) promoter, and an IL-2 signal peptide sequence for secretion of the fusion protein. To express MDK-703, a synthetic DNA fragment was inserted into the vector to fuse MDK1472 to the C-terminus of the IgG2 Fc chain with a (Gly-Ser)_10_ linker between the peptide and the Fc domain. Fusion proteins were transiently expressed in 293 human embryonic kidney cells (FreeStyle® 293-F) by transfecting plasmid DNA into the cells using polyethyleneimine reagent PEI MAX (Polysciences, Inc.). Transfected cells were grown in FreeStyle® 293 Expression Medium (ThermoFisher) in shaker flasks in a 37°C humidified CO_2_ incubator on an orbital shaker rotating at 130 rpm. Cultures were harvested 96 h post-transfection by centrifugation, and the secreted fusion proteins were purified from the supernatants using MabSelect SuRe Protein A affinity columns (Cytiva). The columns were washed with phosphate buffered saline (PBS) and the retained Fc-peptide fusion was eluted with 0.1 M glycine buffer (pH 2.8). Eluates were neutralized with 1 M Tris buffer and quantified by measuring absorbance at 280 nm with a NanoDrop® spectrophotometer. Protein concentrations were determined using calculated extinction coefficients derived from the primary sequence of the protein. Size exclusion chromatography (Superdex 200 Increase) was used to remove high molecular weight impurities prior to measuring the activities of the fusion proteins in bioassays.

### Receptor subunit ECD direct binding and competition binding assays

#### Binding determinations by ELISA

Binding was determined using a competition binding ELISA. Microtiter plates (Immulon 4HBX flat bottom 96-well) were coated with 50 µL of 1 mg/mL human ECDs: IL-7Rα(aa 21-236)-His_6_, Sino Biological Cat# 10975-H08H; cynomolgus IL-7Rα(aa 21-235)-His_6_, Sino Biological Cat# 90332-C08H; human IL-2Rα(aa 23-254)-Fc, Acro Cat# ILG-H5256; cynomolgus IL-2Rα(aa 23-254)-Fc, made in-house, in PBS overnight at 4°C. The wells were emptied and blocked with 300 µL of PBS containing 1% BSA (Bovine Serum Albumin Fraction V, VWR Cat# 97061-416) for 1 h at 25°C. Blocked plates were washed 3 times with 300 µL wash buffer (PBS containing 0.05% Tween-20). C-terminal biotinylated versions of the IL-7R binding peptideMDK1248 and the γc binding peptide, MDK1188 were each pre-complexed with Neutravidin-HRP for at least 45 min prior to use in the assays (10 µM bn-peptide and 0.13 mg/mL NA-HRP in PBS). The test compounds were serially diluted in assay buffer at twice the final desired concentration, 50 µL was added to the assay plate and incubated for 1 h at 4°C. Without washing, 50 µL of pre-complexed biotinylated peptide NA-HRP tracer (at final concentration approximately equal to EC_75_) was added, and the plate returned to 4°C for 45 min. Plates were washed with cold wash buffer, and bound tracer was detected using TMB One Component HRP substrate and Stop Solution (Surmodics Cat# TMBW-1000-01 and LSTP-0100-01). Absorbance at 450 nm was read using Agilent BioTek Synergy HTX Multimode Reader.

#### Binding kinetics determination by biolayer interferometry

Label-free measurement of the equilibrium dissociation constant (K_D_) for peptide binding to IL-7Rα and γc extracellular domains with peptides was done by biolayer interferometry (BLI, Gator^TM^ instrument; Gator Bio, Palo Alto, CA) with small/molecule/antibody/protein (SMAP) biosensors. SMAP surface chemistry allows high-capacity immobilization of biotinylated targets and the measurement of interactions of low molecular weight compounds. BLI experiments were run in 96-well black microtiter plates (200 μL volume per well) at 30°C and a shaker speed of 1000 RPM. Biosensors were preloaded with biotinylated targets or biotin in BLI buffer (PBS containing 0.02% BSA and 0.002% Tween-20). Biotinylated recombinant human receptor extracellular domains (ECD) were purchased from Acro Biosystems IL-7Rα (aa 21-236)-Fc huIgG1 receptor, Cat# IL7-H5258 and hexa-His tagged IL-2Rγ (aa 23-254), Cat# ILG-H85E8). Biotin was purchased from Sigma Cat# B4501. Binding assays were performed as follows: (1) biosensors were preloaded at a high density (7-10 nM) with biotinylated receptor ECDs or biotin (saturating level); (2) 30 sec baseline; (3) 120 sec association for IL-7Rα, and 420 sec for IL-2Rγ; (4) 180 sec for IL-7Rα and 660 sec dissociation for IL-2Rγ; (5) 30 sec baseline. A double-reference procedure was used, wherein two biosensors were loaded with each biotinylated receptor, and two biosensors were loaded with biotin. Receptor coated tips were exposed to increasing peptide concentrations (2-fold dilutions) or no peptide. The second receptor coated tip was iteratively exposed to assay buffer. The negative control tips, coated with biotin, were iteratively exposed to the same peptide titration and buffer as was the receptor of interest. Raw data was processed with the instrument data analysis software (v1.7.2). Double reference subtracted data was processed with the recommended software options and subjected to a Savitzky-Golay filter and fit to a 1:1 binding model (local for IL-7Rα and global for IL-2Rγ).

The curve fit was used to determine the association rate constants (k_a_; units M^-1^ sec^-1^) and the dissociation rate constants (k_d_; units sec^-1^). Equilibrium dissociation constants (K_D_; units nanomolar (nM)) were calculated as a ratio of k_d_/k_a_.

### Preparation of cell lines, PBMCs, and isolated T-lymphocyte subsets

#### Construction and maintenance of cell lines responsive to individual γc cytokines or TSLP

To construct cell lines responsive to each of the γc family cytokines, TF-1 cells were purchased from American Type Culture Collection (ATCC Cat # CRL-2003) and cultured in RPMI 1640 supplemented with 4.5 g/L glucose, 10% FBS, 2 mM L-glutamine, 1mM sodium pyruvate, 10 mM HEPES, penicillin-streptomycin, and 2 ng/mL GM-CSF (TF-1 cells naturally express γc and IL-4Rα). Transfections were performed with an Amaxa Nucleofector device: 2×10^6^ cells were resuspended in 100 μL Nucleofector solution T (Lonza) and added to 2 µg of pcDNA3.1 expression vector encoding full-length IL-2Rβ, IL-7Rα, IL-9Rα, or IL-21Rα; and electroporation was performed with Nucleofector program T-001 (for UniProt accession numbers see S2 Table). Following electroporation, cells were immediately added to RPMI medium and allowed to recover overnight. The next day Geneticin was added to the cultures to select for stable pools of transfected cells. Following antibiotic selection, cells were transitioned from GM-CSF-supplemented medium to medium supplemented with 40 ng/mL IL-2, 15 ng/mL IL-7, 10 ng/mL IL-9, or 25 ng/mL IL-21, respectively.

To construct a TSLP-responsive cell line, the TF-1-7Rα cell line described above was transfected with 2.0 µg pcDNA3.1 expression vector encoding full-length TSLPR (CFLR2, S2 Table), using the Amaxa Nucleofector device with Nucleofector program T-001. Following transfection, the cells were immediately added to RPMI 1640 medium supplemented with 4.5 g/L glucose, 10% FBS, 2 mM L-glutamine, 1 mM sodium pyruvate, 10 mM HEPES, 100 IU/mL penicillin, 100 µg/mL streptomycin, and 20 ng/mL IL-7 and allowed to recover overnight. Hygromycin selection was used to establish a stable pool of cells expressing TSLPR. The cells were cultured for 1-2 weeks in IL-7 supplemented medium and tested by flow cytometry for TSLPR and IL-7Rα expression. The TF-1-7Rα/TSLPR positive pool was then bulk sorted to enrich the high-expressing TSLPR population by Stanford Medicine Shared FACS Facility staff. The sorted pool was cultured in TF-1 RPMI medium supplemented with 20 ng/mL IL-7.

#### Preparation of human PBMCs and lymphocyte subtypes

Human donor buffy coats were purchased from Stanford Blood Center, and PBMCs were isolated by centrifugation in Lymphoprep™ density gradient medium (Stemcell Technologies Cat# 07811). The plasma layer was discarded, the white mononuclear cell layer was decanted and washed twice with cold PBS + 2% FBS, and cell count and viability were determined. For preparation of lymphocyte subsets, naïve CD4+ T-cells were isolated via two-stage negative selection with EasySep™ Human Naïve CD4+ T-Cell Isolation Kit II (Stemcell Technologies Cat# 17555); and CD8+ T-cells were isolated by negative selection from PBMCs with Stem Cell Technology Easy Sep CD8+ kit, or obtained freshly prepared from Stanford Blood Center. Rested naïve CD4+ T-cells or CD8+ T-cells were prepared by overnight incubation at 37°C, 5% CO_2_ in CTS™ OpTmizer™ T-Cell Expansion SFM (ThermoFisher Scientific Cat# A1048501).

### Cell-based assays

#### STAT phosphorylation assays

Cells were deprived of growth factors by overnight incubation at 5×10^5^ cells/mL in starvation medium (RPMI 1640 + 4.5 g/L glucose + 0.2% FBS + 2 mM L-glutamine + 1 mM NaPyr + 10 mM HEPES + 0.5x penicillin-streptomycin with no cytokine supplement) in T75 flasks. For assays, cells were plated in 96-well V-bottom plates at 2×10^5^ cells/well. Three-fold serial dilutions in starvation medium of the test compounds or cytokine were added to the cells and incubated for 30 min at 37°C. Following incubation, cell extracts were prepared by adding a mixture of 10x Cell Lysis Buffer (Cell Signaling Technology Cat# 9803) and 100x HALT Phosphatase and Protease Inhibitor Cocktail (Thermo Fisher Cat# 78442) directly to the wells.

The plates were agitated at 25°C for 5 min to prepare cell extracts for immediate use or to be stored at -80°C until assay. Detection of pSTAT5 was performed using a PathScan® Phospho-Stat5 (pTyr694) Sandwich ELISA Kit (Cell Signaling Technology Cat# 7113). Detection of pSTAT6 was performed using a PathScan® Phospho-Stat6 (pTyr641) Sandwich ELISA Kit (Cell Signaling Technology Cat# 7275). Detection of pSTAT3 was performed using a PathScan® Phospho-Stat3 (pTyr705) Sandwich ELISA Kit (Cell Signaling Technology Cat# 7300). Cell lysates were added to pre-coated ELISA wells and incubated at 4°C overnight. Primary and secondary antibody incubations and wash steps were performed according to the protocols supplied by the manufacturer. Assays were developed using TMB substrate (Surmodics Cat# TMBW-1000-01) for 5-10 min at 25°C and the reaction was stopped using STOP solution (Surmodics Cat# LSTP-1000-01).

To detect phopho-STAT5 in immune subsets of PBMCs by flow cytometry, frozen human and cynomolgus macaques PBMCs (Stemcell Technologies, Cat# 70025.2 and iQ Biosciences, Cat# IQB-MnPB102, respectively) from 5 different healthy donors were thawed and rested overnight in T-cell media (CTS™ OpTmizer™ T-Cell Expansion SFM, ThermoFisher Scientific Cat# A1048501) at 37°C. Cells were stained with a viability dye for 30 min at 37°C followed by surface marker antibodies for 30 min on ice (S3 Table), then treated with serially diluted test compounds in the medium for 30 min. The cells were then washed, fixed, permeabilized as instructed by the manufacturer, and stained with pSTAT5 and intracellular antibodies for 50 min on ice. Antibody-stained cells were analyzed immediately by flow cytometry using the Novocyte Advanteon instrument (Agilent), and the data were analyzed using FlowJo software. Fluorescence minus one (FMO) controls were used to draw gates. For measuring pSTAT5 in activated PBMCs, the overnight rested cells were incubated with CD3/CD28 Dynabeads at 1 bead to 1 cell ratio for 3 days and then rested for 2 days in the fresh medium without the Dynabeads before testing the compounds.

#### Proliferation of PBMC subpopulations

Human PBMCs were isolated from buffy coats by density gradient centrifugation (Lymphoprep®, Stemcell Technologies Cat# 07811) and cultured overnight in T-cell medium (CTS OpTmizer®, ThermoFisher Cat# A1048501) at 3×10^6^ cells/mL in T75 flasks. The following day, cells were resuspended in fresh medium and split into activated and rested groups upon plating. To activate cells, 96-well plates were pre-coated with 10 ng/mL anti-CD3 mAb at 4°C overnight (Purified NA/LE mouse anti-human CD3, BD Cat# 557052). Three-fold serial dilutions of either IL-7 or a test compound were added to the cells and incubated for 4 days at 37°C. After treatment, cells were incubated in viability dye (Live/Dead® Fixable Aqua Cell Stain Kit, ThermoFisher Cat# L34965) for 30 min at 37°C, then stained with surface marker antibodies in PBS + 2% FBS for 30 min on ice. Cells were fixed and permeabilized with Fixation/Permeabilization Buffer (eBioscience Foxp3/Transcription Staining Buffer Set, ThermoFisher Cat# 00-5523-00) for 30 min on ice. Intracellular (Ki-67) staining was then performed in Permeabilization Buffer for 30 min on ice, and the treated cells resuspended in PBS + 2% FBS prior to flow cytometer analysis.

#### IL-7Rα (CD127) internalization assay

CD8+ cells isolated by negative selection from PBMCs with Stem Cell Technology Easy Sep CD8+ kit, or obtained freshly prepared from Stanford Blood Center, were rested overnight in RPMI 1640 containing Pen/Strep, L-Glutamine and 20% FBS; then resuspended to 7.0×10^6^ cells/mL. Surface IL-7R was preloaded by treating the cells with saturating concentrations of the test and control compounds for 20 min on ice, then shifted to 37°C to allow internalization. At various times samples were taken, and each time point was quenched by returning samples to internalization-restrictive ice-cold conditions. Following collection of the timed incubations, the cells were washed 2x with cold PBS + 2% FBS and stained with anti-CD127-Alexa Fluor® 488 (anti-IL-7Rα) (antibody clone 40131; R&D Systems Cat# FAB306G) for 30 min on ice, protected from light, and fixed with 100 μL of 1.5% paraformaldehyde (16% stock diluted into PBS). To determine levels of cell surface IL-7Rα, analysis was performed by flow cytometry to determine median fluorescence intensity (MFI) of cells at each condition.

Negative controls consisted of a blank (no compound added); MDK1169, a peptide agonist of another γc family receptor (which contains a γc binding domain similar to MDK1472 but lacking the IL-7Rα binding domain); and MDK-202, an Fc-fusion of MDK1169 (a construct analogous to MDK-703, with the same IgG2 Fc domain fusion partner) [59]. A parallel assay with all compounds held on ice for the duration of the time course was also performed. Flow data was collected as MFI, and for display, data was normalized with blank value setting the upper limit of 100% surface CD127, and the signal baseline (0%) estimated by adding an excess of unlabeled antibody clone 40131 for 30 min on ice prior to staining with the labeled detection antibody.

#### Analysis of effects of test compounds on activation of all gamma common family receptors, and the TSLP receptor

TF-1-derived lines separately expressing each gamma common family private receptor were starved overnight at 5×10^5^ cells/mL. The following day, the cells were counted and plated at 2×10^6^ cells/well in 96-well plates. Dose response assays of pSTAT activation by the gamma common cytokines IL-2, IL-4, IL-7, IL-9, IL-15, and IL-21 were performed with three-fold serial dilutions of the cytokines added to the respective cytokine receptor-expressing TF-1 cells and incubated for 30 min at 37°C to estimate EC_75_ levels. To assess inhibition of the activities of these cytokines, the test compounds MDK1188, MDK1248, MDK1472, or MDK-703 were serially diluted in starvation medium, and pre-incubated with the cells for 20 min at 37°C. Sub-maximal levels (EC_75_) of each cytokine were then added to the corresponding cells for 30 min at 37°C. Cell extracts were prepared and analyzed for pSTAT3, pSTAT5, or pSTAT6 accumulation with Pathscan pSTAT ELISA kits as described above. Human IL-2, IL-4, IL-7, IL-9, IL-15, and IL-21 were obtained from Peprotech (Cat#s 200-02, 200-04, 200-07, 200-09, 200-15, and 200-21).

To evaluate the effect of MDK-703 on TSLP receptor-mediated signaling, pSTAT5 assays were run using the engineered IL-7Rα/TSLPR cell line. In each assay, the cells were starved overnight at 5×10^5^ cells/mL, counted, and 1.8×10^5^ cells/well were plated in 96-well plates. These cells respond to both IL-7 and to TSLP with the production of pSTAT5. To separately measure TSLP receptor activation in the presence of the IL-7R agonist MDK-703, the interaction of MDK-703 with γc was blocked by the inclusion of 20uM of a high affinity competitive inhibitor of the interaction, MDK2058. IL-7 and TSLP receptor activation assays were run 30 min at 37^°^C and pSTAT5 accumulation was detected with PathScan Phospho-Stat5 (Tyr694) Sandwich ELISA Kit, Cell Signaling Technology Cat#7113 to quantitate the response. Absorbance at 450 nm was read using Agilent BioTek Synergy HTX Multimode Reader (Figs 2 A and 2B). In the same assay, the dose response of pSTAT5 activation by TSLP was measured with or without preincubation with the gamma common chain blocker MDK2058, +/-MDK-703 at EC_95_ (600 nM final concentration), for 4 min at 4°C (Fig 4H).

#### PBMC proliferation

Frozen human PBMCs from 5 healthy donors (Stemcell Technologies, Cat# 70025.2, Lot numbers 2108409012, 2109407009, 210680104C, 210680801C, 211280501C, 220781802C) were thawed and rested overnight at 37°C, 5% CO_2_ in CTS™ OpTmizer™ T-Cell Expansion SFM (ThermoFisher Scientific Cat# A1048501). Four million cells in 2 mL media were cultured in the plate coated with or without 10 ng/mL of CD3 antibody (Clone SP34-2) and in the presence or absence of 100 nM MDK-703 or 1 nM IL-7. Fresh media containing MDK-703 and IL-7 were provided every 4-5 days. At indicated time points, cell aliquots were taken, and cell surface staining was performed using antibodies shown in S3 Table followed by viability dye staining. For Ki-67 and FoxP3 staining, surfaced stained cells were fixed and permeabilized according to the manufacturer’s instructions and analyzed immediately using the Novocyte Advanteon instrument (Agilent). The data were analyzed using FlowJo software. Fluorescence Minus One (FMO) controls were used to draw gates. Percentages of Ki-67+ cell populations were expressed as average ± SEM. Memory T-cells are as follows: Tn (CD45RA+CCR7+CD28+CD95-), Tscm (CD45RA+CCR7+CD28+CD95+), Tem (CD45RA-CCR7-), Tcm (CD45RA-CCR7+), and Temra (CD45RA+CCR7-). To count cells per well, at each assay time point 50 μL of the cells were taken from each well and incubated with 200 μL of the viability dye solution. The NovoCyte Advanteon flow cytometer was set to analyze precisely 50 μL (12.5 μL original culture volume), which allowed the calculation of the absolute live cell counts. Because the culture volumes change over time due to splitting and removal, dilution factors were considered when calculating the total number of cells.

#### Pharmacodynamic study with humanized NSG mice

The PD study was performed at Crown Biosciences (San Diego, CA). Ten HIS-NSG mice (NSG™ NOD.Cg-Prkdcscid Il2rgtm1Wjl, Stock No. 005557) engrafted with CD34+ cells from 2 donors (The Jackson Laboratory, Bar Harbor, ME) were treated intravenously once with 1 mg/kg of MDK-703 or a human Fc control. Blood samples were collected via submandibular bleeding on day 7, and terminal blood samples were collected via cardiac puncture. Red blood cells were removed using RBC Lysis Buffer (BioLegend Cat#-420302). Spleens were harvested on day 12 and processed to single-cell suspensions using the Gentle MACS, program m_spleen_04. Cell aggregates were removed by passing through a 70 μm pore filter, and red blood cells were lysed. Splenocytes were incubated with TruStain FcX before antibody staining as described above. Antibody-stained cells were acquired by using BD LSRFortessa X-20 flow cytometer and analyzed with Kaluza software.

#### Single dose pharmacokinetic and pharmacodynamic studies of MDK-703 in NHP

NHP studies were conducted at Envol Biomedical (Naples, FL). For the PD study, 3 female cynomolgus macaques were dosed once subcutaneously at 0.3 mg/kg MDK-703, and blood samples were collected at days 1(pre-dose), 2, 3, 4, 6, 8, 12, 18, 24, 30, 36, and 42 days and analyzed by complete blood count and flow cytometry. For the PK study, 3 female macaques per group were dosed once with 1 mg/kg MDK-703 via intravenous bolus, subcutaneous, or intramuscular injection. Blood samples were collected at Pre-Dose (day 1), 0.5, 1 h, 2 h, 4 h, 8 h, 24 h, 48 h, 72 h, 96 h, 120 h, 144 h, 168 h, 192 h, and 216 h post-dose into serum separator vials and processed for serum (30 min at room temperature followed by centrifugation at 4°C, 3000 x g for 5 min). The serum was transferred to a polypropylene tube and stored at -80°C until analysis. A standard sandwich ELISA using a three-fold dilution of serum samples was performed to quantify the amount of MDK-703. For plasma samples from subcutaneously injected animals, a biotinylated rat polyclonal antibody against human Fc was used as a capture antibody and anti-human IgG Fc-HRP (GenScript Cat# A01854-200) as a detection antibody. For plasma samples from the intravenous or intramuscular injection, biotinylated mouse anti-human IgG2 Fc (Southern Biotech Cat# 9070-08) was used as a capture antibody and rabbit polyclonal antibody raised against IL-7Rγ peptide agonist followed by a mouse anti-rabbit IgG Fc HRP (Genscript Cat# A01856-200) as a detection antibody. The quantity of MDK-703 in each serum sample was calculated with a standard curve of the test compound diluted in the control cynomolgus monkey serum. Individual animal serum MDK-703 concentration versus time data was imported into Phoenix WinNonlin v8.3 (Certara, Princeton, NJ) for analysis. All values that were less than the lower limit of quantitation (LLOQ) of the assay were treated as missing for the PK analyses. The individual animal concentration versus time data was analyzed using a noncompartmental model with extravascular administration. Nominal dose and sample collection times were used in estimating the PK parameters.

## Acknowledgments

The authors would like to acknowledge Julie Crider, PhD, for medical editing contributions.

