## Supplemental Material for "A mechanistically novel peptide agonist of the IL-7 receptor that addresses limitations of IL-7 cytokine therapy"

### Supporting Material

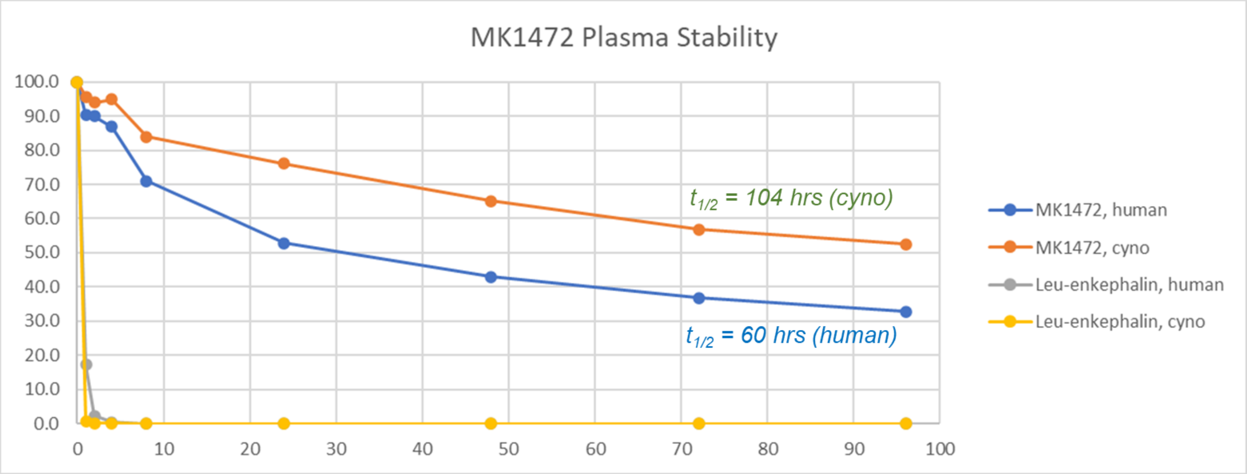

**S1 Fig. Plasma stability of MDK 1472 *in vitro*.** 10 mM MDK1472 or control peptide YGGFL incubated in duplicate in human or cynomolgus plasma at 37°C. At indicated time points, compounds were isolated from the plasma and the quantity of intact peptides measured by LC/MS/MS (Study performed by Quintara Discovery, Hayward, CA).

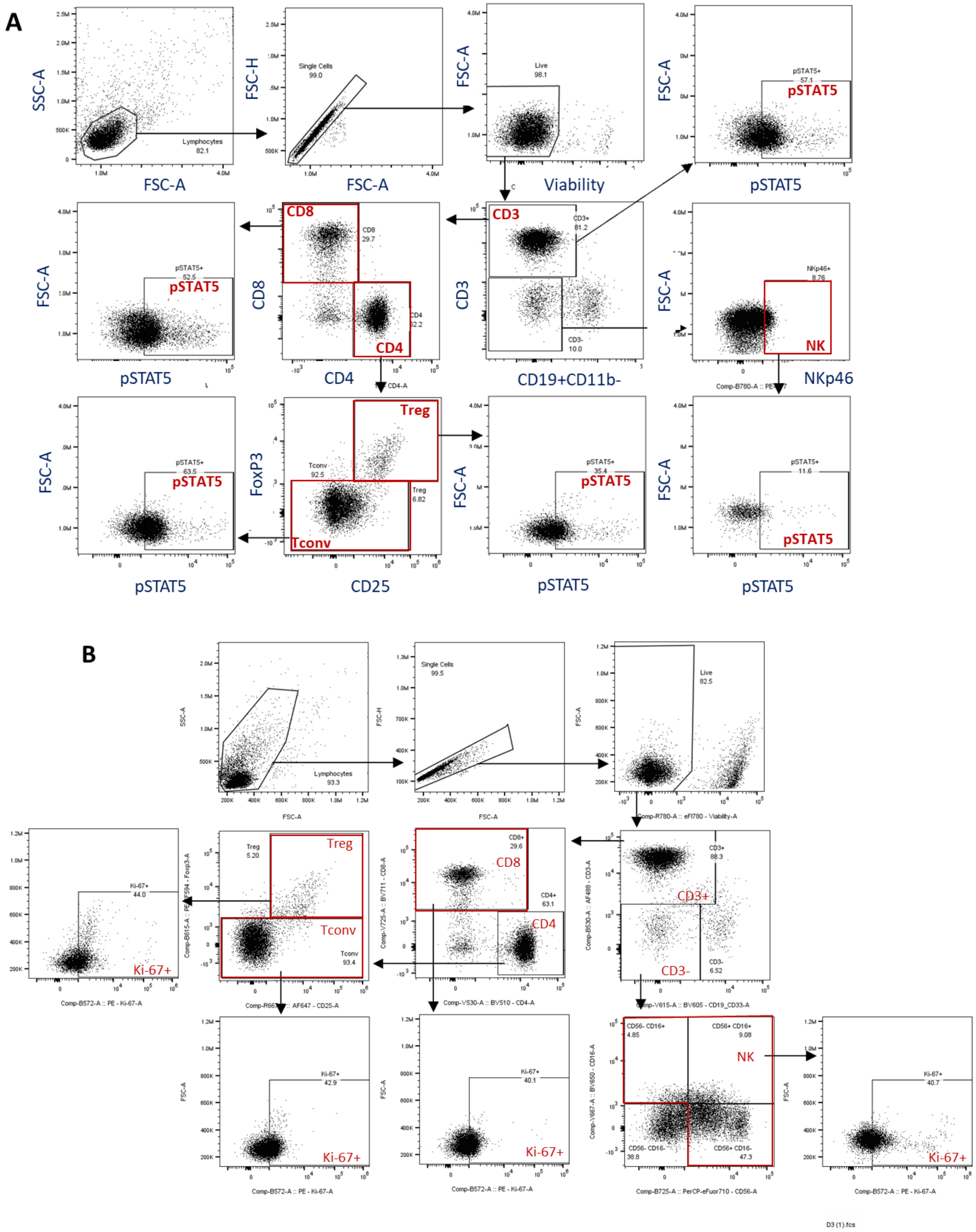

**S2 Fig. Flow cytometry gating scheme for STAT5 and Ki-67 determinations in major lymphocyte populations of human PBMCs. (A) pSTAT5 assay. (B) Ki-67 assay.**

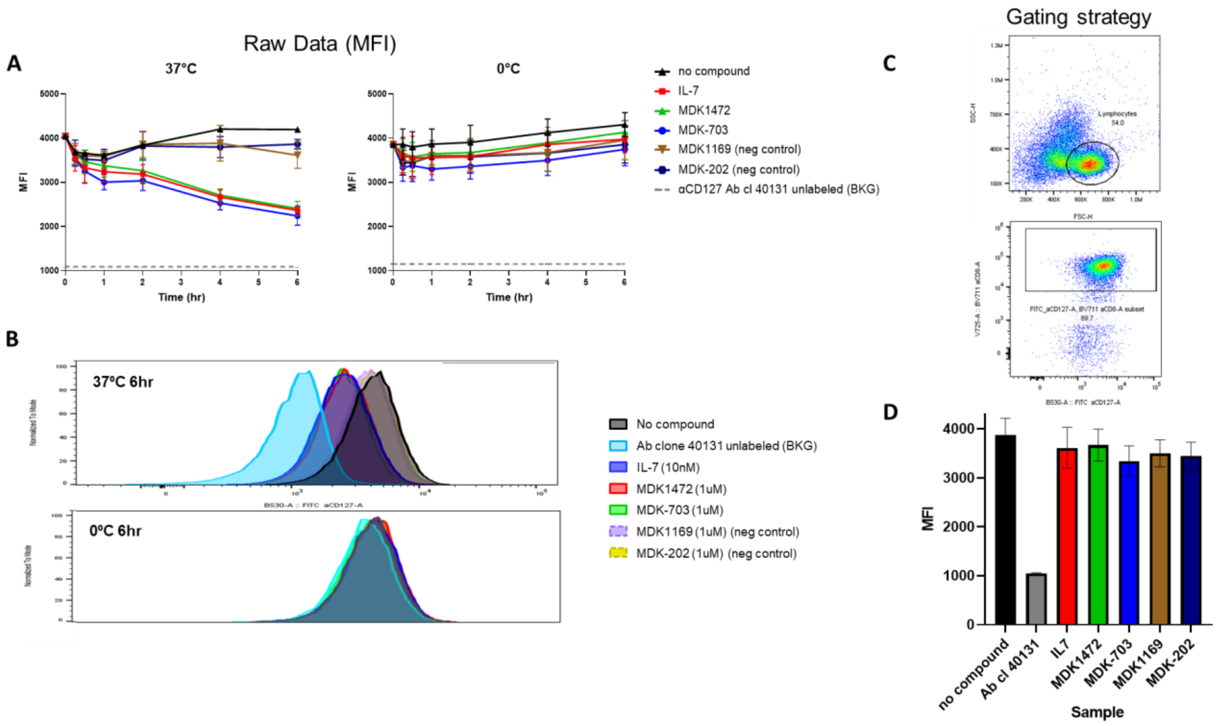

**S3 Fig. Flow cytometry study of IL-7Rα internalization +/- exposure to test compounds.**

Primary MFI data (A) plotted time course, (B) histograms, (C) gating scheme. (D) Evaluation of interference of IL-7 and IL-7Rα agonist test compounds, and irrelevant test compounds with detection of cell surface IL-7Rα on cell surface by labelled anti-CD127 Ab clone #40131. All test compounds are loaded on the cells on ice for 35 min, stained, and analyzed by flow (as described in detail in Methods).

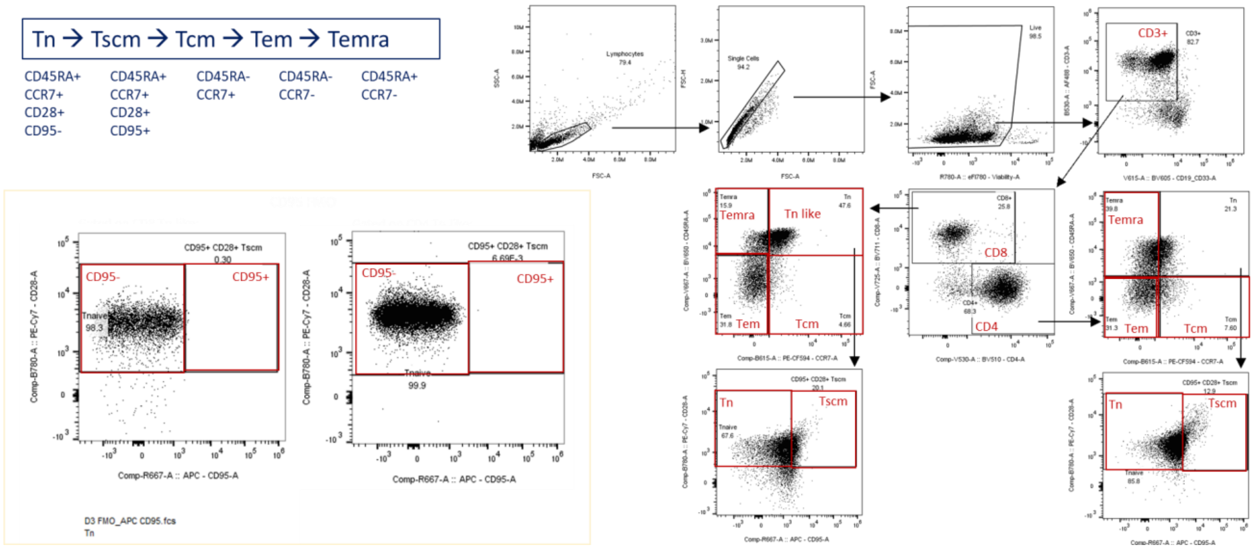

**S4 Fig. Flow cytometry gating scheme for analysis of T-memory cell Ki-67 expression and expansion of cell numbers following *in vitro* exposure of human PBMC to test compounds.**

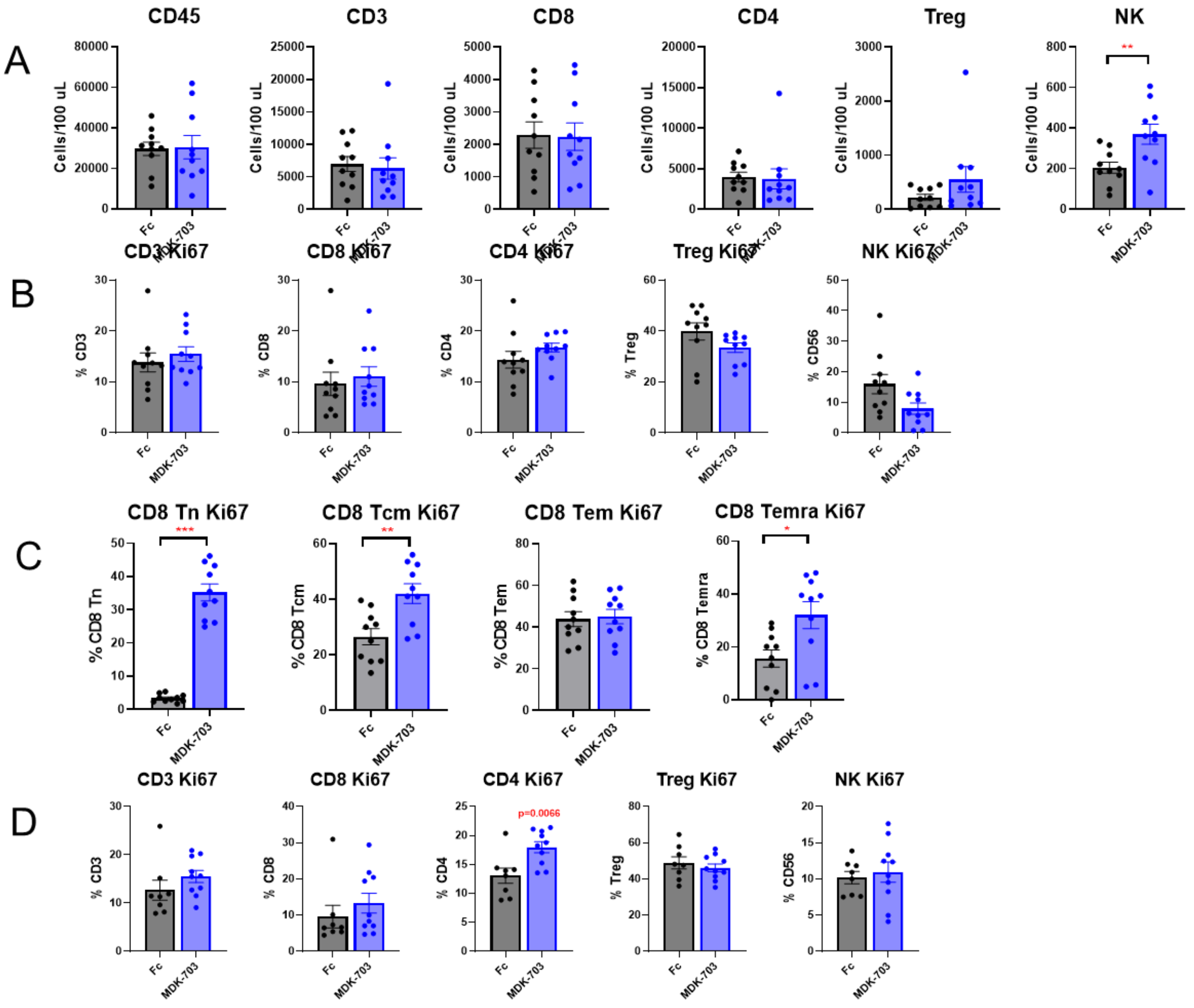

**S5 Fig. Proliferation of major lymphocyte subpopulations in humanized mice treated with test compounds.** (A) Day 7 cell counts in blood. (B) Day 12 Ki-67 in blood. (C) Day 7 memory T-cells Ki-67 in blood. (D) Day 12 Ki-67 in spleen**.**

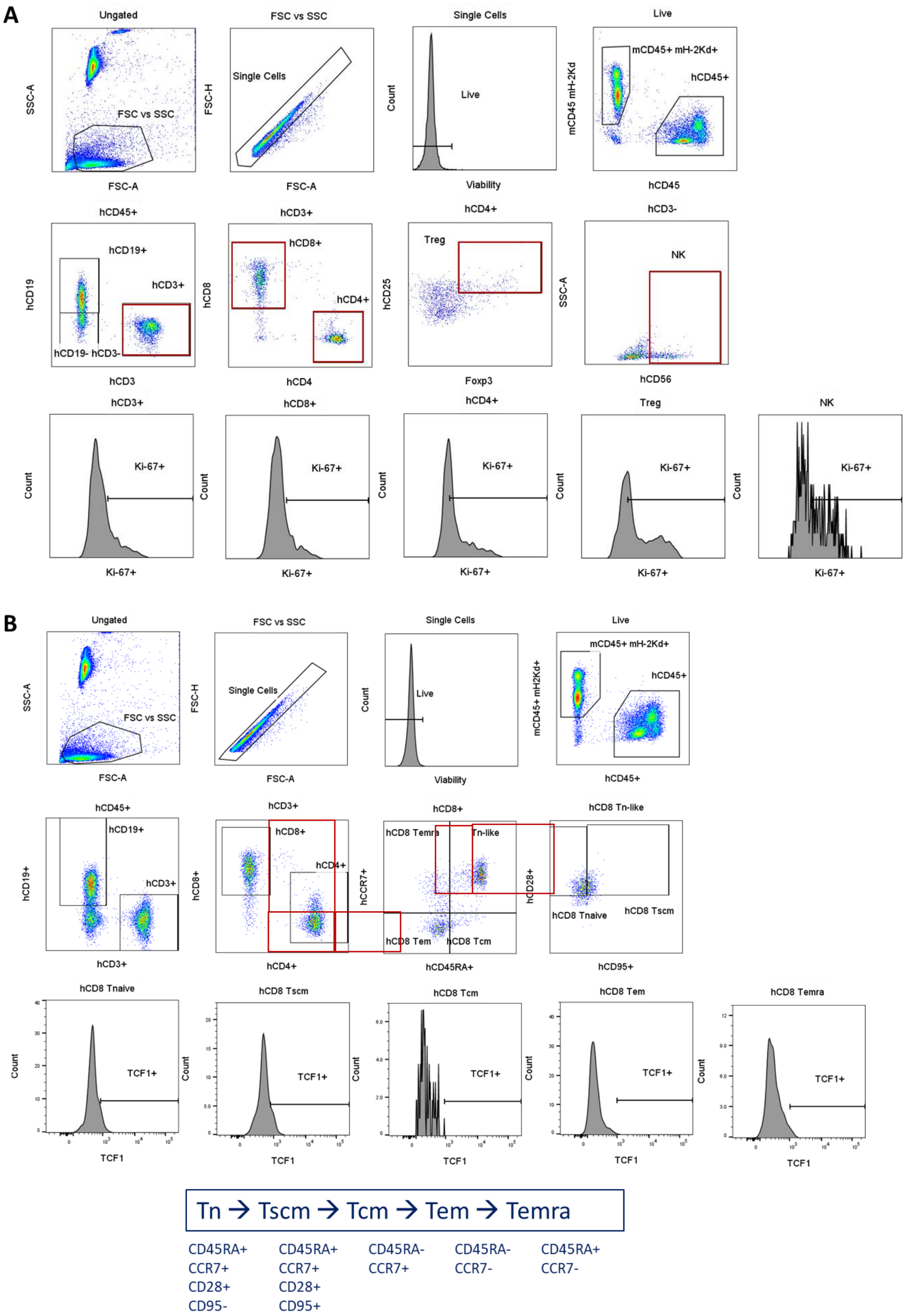

**S6 Fig. HIS mouse flow cytometry sample gating information.** (A) TBNK Ki-67+. (B) Tn and Tmem TCF1+.

#### S1 table. Potency values

| Human | | EC50 | | | |
|---|---|---|---|---|---|
| | | **CD8** | **CD4** | **Treg** | **NK** |
| **Rest** | **IL-7** | 4.4E-12 | 4.4E-12 | 6.3E-12 | N/A |
| | **MDK-703** | 7.2E-10 | 7.8E-10 | 1.6E-09 | N/A |
| **CD3/CD28** | **IL-7** | 8.7E-12 | 8.8E-12 | 2.5E-11 | N/A |
| | **MDK-703** | 2.7E-10 | 2.6E-10 | 1.1E-09 | N/A |
| NHP Rest | **IL-7** | 2.4E-12 | 1.2E-12 | 5.1E-12 | N/A |
| | **MDK-703** | 1.4E-09 | 8.3E-10 | 6.5E-09 | N/A |

#### S2 Table. UniProt accession numbers.

| **Protein Name** | **UniProt Accession Number** |
|---|---|
| Human IgG2 Fc (MDK-703 expression construct) | P01859 |
| Human IL-2Rβ | P14784 |
| Human IL-4Rα | P24394 |
| Human IL-7Rα | P16871 |
| Human IL-9Rα | Q01113 |
| Human IL-15Rα | Q13261 |
| Human IL-21Rα | Q9HBE5 |
| Human CRLF2 transcript variant 1 Gene ORF cDNA clone expression plasmid (TSLPR) | Q9HC73 |
| Human IL-2 | P60568 |
| Human IL-4 | P05112 |
| Human IL-7 | P13232 |
| Human IL-9 | P15248 |
| Human IL-15 | P40933 |
| Human IL-21 | Q9HBE4 |
| Cynomolgus IL7Rα | Q38IC7 |
| Cynomolgus IL-7 | A0A2K5W745 |

### S3 Table. Labelled antibody reagents for flow cytometry

| Antibody | Fluorochrome | Clone | Company | Catalog number |
|---|---|---|---|---|
| **CCR7** | PE-CF594 | 150503 | BD Biosciences | 562381 |
| **CCR7** | PE/Cy7 | G043H7 | BioLegend | 353226 |
| **CD3** | AF488 | SP34.2 | BD Biosciences | 557705 |
| **CD3** | AF700 | SP34.2 | BD Biosciences | 557917 |
| **CD3** | BUV395 | SK7 | BD Biosciences | 564001 |
| **CD4** | BV510 | SK3 | BD Biosciences | 562970 |
| **CD4** | BV650 | L200 | BD Biosciences | 563737 |
| **CD4** | APC | SK3 | BioLegend | 344614 |
| **CD8** | BV711 | SK1 | BioLegend | 344734 |
| **CD8** | BUV737 | SK1 | BD Biosciences | 612755 |
| **CD8** | PE | BW135/80 | Miltenyi | 130-113-158 |
| **CD8** | PE-Cy7 | SK1 | BioLegend | 344712 |
| **CD8** | PE-eFluor 610 | RPA-T8 | eBioscience | 61-0088-42 |
| **CD11b** | BV605 | M1/70 | BioLegend | 101257 |
| **CD127** | PE | 40131 | R&D Systems | FAB306P |
| **CD159a** | PE-Cy7 | BAB281 | Beckman Coulter | A60797 |
| **CD16** | BV650 | B73.1 | BD Biosciences | 740639 |
| **CD19** | BV605 | HIB19 | BioLegend | 302244 |
| **CD19** | BV711 | HIB19 | BioLegend | 302246 |
| **CD20** | PE-Vio770 | LT20 | Miltenyi | 130-113-375 |
| **CD25** | AF647 | M-A251 | BioLegend | 356128 |
| **CD25** | AF780 | CD25-4E3 | ThermoFisher | 47-0257-42 |
| **CD25** | BB700 | M-A251 | BD Biosciences | 566447 |
| **CD25** | APC-eFluor780 | CD25-4E3 | Invitrogen | 47-0257-42 |
| **CD25** | BV510 | M-A251 | BioLegend | 356120 |

| | | | | |
|---|---|---|---|---|
| **CD27** | PerCP-Cy5.5 | O323 | BioLegend | 302820 |
| **CD28** | PE-Cy7 | CD28.2 | BioLegend | 302926 |
| **CD33** | BV605 | P67.6 | BioLegend | 366612 |
| **CD45** | AF488 | HI30 | BioLegend | 304017 |
| **CD45** | BV785 | HI30 | BioLegend | 304048 |
| **CD45RA** | BV650 | HI100 | BioLegend | 304135 |
| **CD45RO** | BV421 | UCHL1 | BioLegend | 304224 |
| **CD56** | PerCP-eFluor 710 | CMSSB | Invitrogen | 46-0567-42 |
| **CD56** | BV605 | HCD56 | BioLegend | 318334 |
| **CD95** | APC | DX2 | BD Biosciences | 558814 |
| **FoxP3** | PE-CF594 | 236A/E7 | BD Biosciences | 563955 |
| **FoxP3** | PE | 206D | BioLegend | 320108 |
| **FoxP3** | PE | PCH101 | Invitrogen | 12-4776-42 |
| **FoxP3** | PE | 259D | eBioscience | 12-4776-42 |
| **Ki67** | PE | SolA15 | Invitrogen | 12-5698-82 |
| **Ki67** | BV421 | B56 | BD Biosciences | 562899 |
| **Ki67** | PerCP/Cy5.5 | Ki-67 | BioLegend | 350520 |
| **Live/Dead** | eFl780 | - | Invitrogen | 65-0865-18 |
| **Live/Dead** | Pacific Orange | - | Invitrogen | L34967 |
| **mCD45** | FITC | 30-F11 | BioLegend | 103108 |
| **mH-2Kd** | FITC | SF1-1.1 | BioLegend | 116605 |
| **NKp46** | PE-Cy7 | BAB281 | Beckman Coulter | B38703 |
| **pSTAT5** | BV421 | 47/Stat5 (pY694) | BD Biosciences | 562984 |
| **TCF-1** | AF488 | S33-966 | BD | 567018 |
